# Bacterial Protein Signatures Identified in *Porphyromonas gingivalis* Containing-Autophagic Vacuoles Reveal Co-Evolution Between Oral Red/Orange Complex Bacteria and Gut Bacteria

**DOI:** 10.1101/2024.07.11.602567

**Authors:** Ayana Paul, Bridgette Wellslager, Maddie Williamson, Özlem Yilmaz

## Abstract

Modern oral bacterial species present as a concoction of commensal and opportunistic pathogens originating from their evolution in humans. Due to the intricate colonization mechanisms shared amongst oral and gut bacteria, these bacteria have likely evolved together to establish and adapt in the human oro-digestive tract, resulting in the transfer of genetic information. Our liquid chromatography-with-tandem-mass-spectrometry (LC-MS-MS) analyses have revealed protein signatures, Elongation Factor Tu, RagB/SusD nutrient uptake outer membrane protein and DnaK, specifically from *Porphyromonas gingivalis*-containing autophagic vacuoles isolated from the infected human primary gingival epithelial cells. Interestingly, our Mass-Spectrometry analysis reported similar proteins from closely related oral bacteria, *Tannerella forsythia* and *Prevotella intermedia*. In our phylogenetic study of these key protein signatures, we have established that pathogenic oral bacteria share extensive relatedness to each other and gut resident bacteria. We show that in the virulence factors identified from gut bacteria, Elongation Factor Tu and DnaK, there are several structural similarities and conservations with proteins from oral pathogenic bacteria. There are also major similarities in the RagB/SusD proteins of oral bacteria to prominent gut bacteria. These findings not only highlight the shared virulence mechanisms amongst oral bacterial pathogens/pathobionts but also gut bacteria and elucidate their co-evolutions in the human host.

## 1. Introduction

A microbiome can be defined as the ecological community of pathogenic and commensal microorganisms that inhabit a particular area of the body. The oral microbiome is likely one of the most influential microbiomes of the human body as the oral cavity is a point of entry of microorganisms into other areas of the body and thus may play a role in the pathologies outside of this area (Willis and Gabaldón, 2020, Dewhirst et al., 2010). This microbiome is substantially diverse and can be divided into several niches including the buccal mucosa, throat, tongue, saliva, gingiva, and sub-gingiva (Lee et al., 2021, Faran Ali and Tanwir, 2012). The modern human oral microbiota is believed to have evolved from changes diet, environment, hygiene, health, genetics, lifestyle over human evolution and industrialization (Shaw et al., 2017). Characterization of the modern oral microbiome has resulted in the organization of microbial complexes of varying association with healthy and diseased states of the oral cavity (Avila et al., 2009). Blue, purple, green and yellow complex bacteria are understood to colonize and proliferate in the oral cavity in the early stages of oral microbiome development. A number of bacteria are commonly found in healthy oral microbiomes such as, *Streptococcus, Staphylococcus, Veillonella* and *Eubacteria* (Avila et al., 2009, Wilson, 2004). Conversely, bacteria that have been characterized as pathogenic and associated with diseased states are *Prevotella* species, *Tannerella forsythia, Treponema denticola* and *Porphyromonas gingivalis* (Haffajee and Socransky, 1994, Spooner et al., 2016, Haffajee and Socransky, 2005). These bacteria, commonly characterized as orange and red complex bacteria, colonize the oral microbiome last and enable the proliferation of other pathogenic bacteria and induction of host proinflammatory responses. Figure 1 briefly demonstrates how the oral bacterial complexes are established and how these oral complexes can lead to disease pathologies and translocation in the human body. Poor oral hygiene and antibiotic use are important contributing factors to oral dysbiosis (Dewhirst et al., 2010, Jiao et al., 2014). Thus, dysbiosis of the oral microbiome can result in numerous oral diseases such as gingivitis, dental carries and periodontitis (Jiao et al., 2014, Simón-Soro et al., 2013). The number of people with oral conditions in general have also increased globally from 1990 to 2015 with disability adjusted life years (DALYS) increasing by 64% (Kassebaum et al., 2017). The organization, pathogenicity and dysbiosis of the oral microbiome are pertinent to our understanding of oral disease outcomes.

**FIGURE 1.**
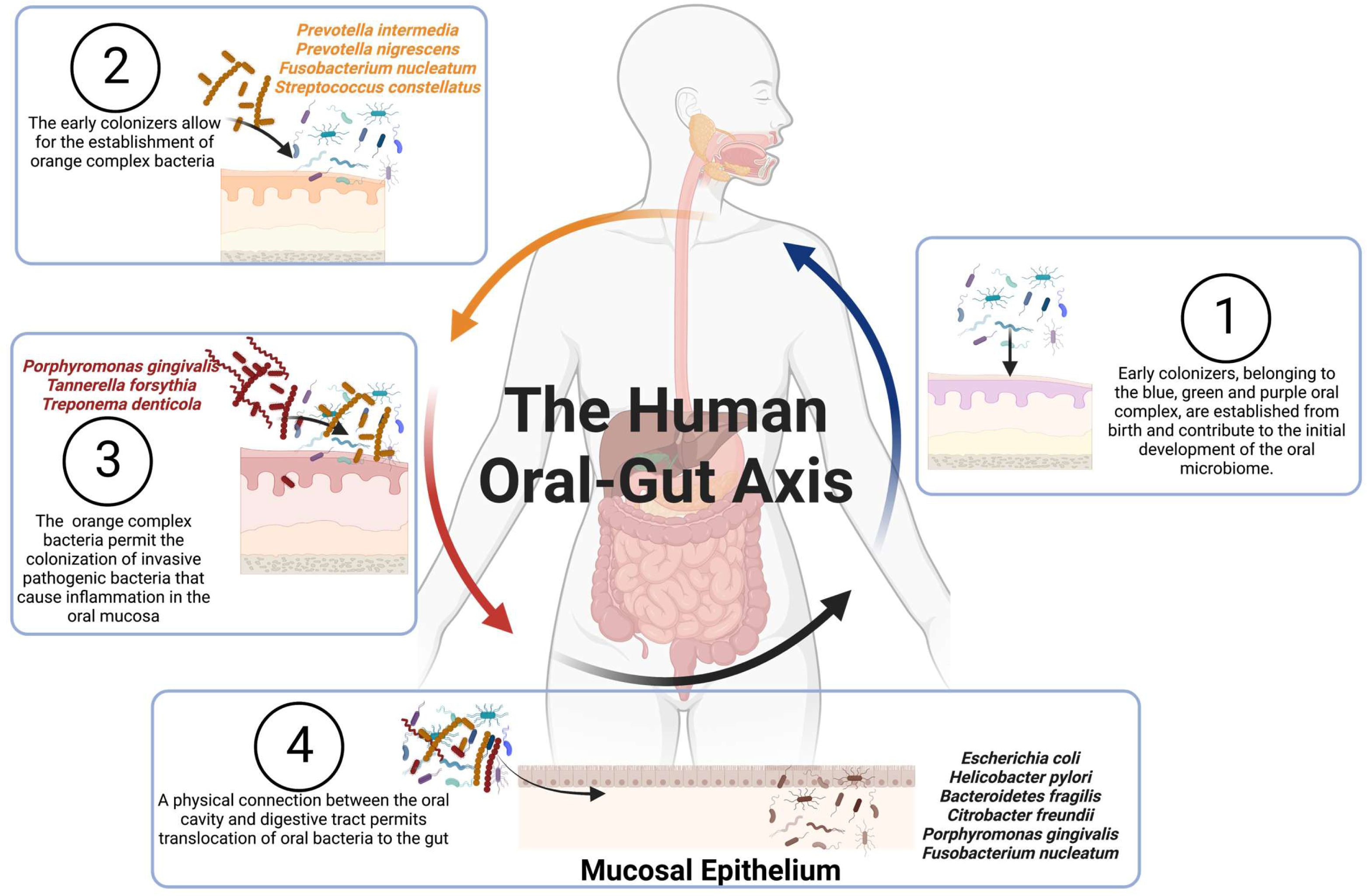
The Human oral-gut axis can be organized into different core complexes that are linked via physical connections. A schematic diagram demonstrating the organization of key bacteria within the oral microbiome and its physical connection to external microbiomes such as the gut microbiome. Bacteria are not limited to their prospective niches and have physical access to different areas of the body.

The oral microbiome is a unique ecological space in that is physically connected to other areas of the body, especially the gut microbiome (Figure 1) (Narengaowa et al., 2021). Due to this accessibility, it is likely that the oral microbiota shares similar evolutionary histories with the gut microbiome. In fact, in a study conducted by The Human Microbiome Project Consortium, it was found 125 bacterial species consistently present in both saliva and stool from 470 people (Human Microbiome Project, 2012). Despite the consistencies in disease associations and physical connections in the microbiomes, the relationship between oral and gut bacteria is incredibly understudied. Studying the co-evolution of these bacteria and their molecular mechanisms could potentially shine a light on novel holistic therapeutic strategies for non-communicable microbially associated diseases.

*P. gingivalis*, is a Gram (-), obligate anaerobic bacterium and is a member of the oral red complex within the oral cavity. This bacterium demonstrates a unique ability to persist and spread within gingival epithelial cells (GECs), which serve as a microbial protective barrier in the oral mucosa (Lee et al., 2020a, Nakayama and Ohara, 2017, Mohanty et al., 2019, Jenkinson and Lamont, 2005, Atanasova et al., 2016, Yilmaz et al., 2006). *P. gingivalis* can avoid the host immune system and induce dysbiosis of the oral microbiome and possesses a number of virulence factors to aid in its pathogenesis such as lipopolysaccharides, gingipains, out membrane vacuoles as well as pili (Jia et al., 2019, Chen et al., 2001a, Zhang et al., 2021). Our lab has begun to characterize the intracellular life of *P. gingivalis* and has revealed that the microorganism can evade a variety of cellular mechanisms designed to eliminate pathogens (Roberts et al., 2017a, Lee et al., 2020b, Yao et al., 2010) (Spooner et al., 2014). One key mechanism that *P. gingivalis* modulates is xenophagy in order to avoid phagolysosomal fusion while inside the Endoplasmic Reticulum (ER)/LC3 rich double-membrane vacuoles for which *P. gingivalis* can replicate and survive in (Lee et al., 2018b, Wellslager et al., 2024). In fact, *P. gingivalis* is most strongly associated with chronic periodontitis, extensive inflammation of the periodontal tissue leading to tooth loss, and demonstrates unique virulence mechanisms for subduing host immune response, positioning as a major pathogen within the oral microbiome (Hajishengallis et al., 2012, Lee et al., 2018a, Lee et al., 2020c, Lee et al., 2020d, Roberts et al., 2017b, Yilmaz, 2008, Lee and Yilmaz, 2020, Atanasova and Yilmaz, 2015, Nakhjiri et al., 2001) Our recent study into the bacterial proteomics involved in the vacoular life of this bacterium using Mass Spectrometry has prompted study of several bacterial protein signatures (Lee et al., 2018b, Wellslager et al., 2024, Chowdhury et al., 2024). Elongation Factor Tu, DnaK and SusD nutrient uptake outer membrane protein signatures, belonging to pathogenic oral bacteria *Prevotella intermedia* and *T. forsythia* were identified inside of *P. gingivalis* containing autophagic vacuoles which were selectively isolated for *P. gingivalis* from the infected primary GECs as we recently described (Wellslager et al., 2024). These bacteria are both closely related *P. gingivalis* and of the identical or proximal oral complex (Mohanty et al., 2019). It is not fully understood what functional role these proteins play in the survival of *P. gingivalis* in the autophagic vacuoles. Further, the evolution of established pathogenic oral complexes and their shared survival mechanisms in host niches is not well understood. Lastly, the extent of relatedness of these protein signatures from *P. gingivalis*, related pathogenic oral and gut bacteria is not known as well. In this study, our objective was to determine the extent of relatedness of proteins identified in *P. gingivalis* contained vacuoles across oral complex bacteria to understand the evolution, maintenance, and establishment of this microbiome and their established bacterial pathogens. We have found that proteins identified from red and orange complex bacteria *T. forsythia* and *P. intermedia* are evolutionary related with bacteria from not only the oral microbiome but the gut microbiome based on phylogenetic analysis. Many of these bacteria have well characterized virulence mechanisms involving the identified protein signatures. The predictive secondary structures of these proteins belonging to these oral pathogens offer additional support for homology and conserved structure. Understanding the evolutionary dynamics of these bacterial species may aid in our understanding of the virulence and survival mechanisms amongst oral bacteria as well as gut bacteria. Extensive study into the protein homology shared across these two microbiomes may contribute to the growing interest in the oral-gut axis and its connection to systemic diseases.

## 2. Materials and Methods

### The Magnetic Isolation of *P. gingivalis-*Containing Autophagic Vacuoles

*P. gingivalis* ATCC 33277 strain was incubated in 5 μg/mL Lipobiotin (LB) in PBS overnight at 4°C. Labelled *P. gingivalis* was then incubated with MagCellect Streptavidin Ferrofluid (R&D Systems) for 20 min at 4°C and was used to infect GECs at MOI 100 for 24 h. *P. gingivalis-*specific autophagic vacuoles were then selectively isolated as we previously described (Wellslager et al., 2024, Chowdhury et al., 2024). In brief, GECs were washed with Buffer A (15 mm HEPES buffer with 20 mm sucrose, 50 mm MgCl_2_, and protease inhibitors containing 0.02% ethylenediaminetetraacetic acid (EDTA)) to remove non-ingested bacteria, were trypsinized, and were resuspended in Buffer A combined with Benzonase Nuclease (25 U/mL; Novagen) and Cytochalasin D (5 µg/mL; Sigma). Following resuspension, *P. gingivalis*-containing autophagic vacuoles were released via repeated sonication on ice via a ThermoFisher Qsonica Sonicator Q500 (FisherSci) at an amplitude of 75% for 10 s, until 90% cellular disruption was visibly noted. GECs were centrifuged at 300 g for 2 min between sonication steps, after which the supernatant was removed and replaced with 500 μL of Buffer A. The samples were then left on ice for 10 min, after which, the supernatants were removed and incubated at 37°C for 5 min. Following incubation, the samples were placed in a Magnetic Rack (Bio-Rad) for 30 min prior to repeated manual rinsing of the magnetic fraction with Buffer A at a flow rate of 250 μL/min for 15 min to ensure purity. Finally, the autophagic vacuole-containing fractions were collected and centrifuged at 15,000 g for 12 min, before the supernatant was decanted and the autophagic vacuoles were resuspended in 100 μL of HEPES buffer combined with protease inhibitors.

### Mass Spectrometry Sample preparation

In preparation for LC-MS/MS Mass spectrometry, *P. gingivalis*-specific autophagic vacuoles were then combined with 9M urea, 50 mM Tris pH 8, and 100 units/mL Pierce Universal Nuclease (ThermoScientific cat. # 88700) buffer to gently release the autophagic vacuoles’ contents. The concentration of protein was then measured utilizing a BCA assay (ThermoScientific cat. # 23225). Proteins were then incubated with LysC and trypsin (Sigma cat. # T6567) at 37°C for 18 hours to promote digestion, and the resulting peptides were desalted using C18 Stage tips (Pierce cat. # 84850).

### Liquid Chromatography and Mass Spectrometry Data Acquisition Parameters

Peptides were separated and analyzed on an EASY nLC 1200 System (ThermoFisher) in-line with the Orbitrap Fusion Lumos Tribrid mass spectrometer (ThermoFisher) with instrument control software v. 4.2.28.14. 2 ug of tryptic peptides were pressure loaded at 1,180 bar and peptides were separated on a C18 reversed phase column (Acclaim PepMap RSLC, 75 µm x 50 cm (C18, 2 µm, 100 Å) ThermoFisher) using a gradient of 5% to 40% B in 180 min (Solvent A: 5% acetonitrile/0.1% formic acid; Solvent B: 80% acetonitrile/ 0.1% formic acid) at a flow rate of 300 nL/min with a column heater set to 50°C.

Mass spectra were acquired in data-dependent mode with a high resolution (60,000) FTMS survey scan, mass range of m/z 375-1575, followed by tandem mass spectra (MS/MS) of the most intense precursors with a cycle time of 3 s. The automatic gain control target value was 4.0e5 for the survey MS scan. Fragmentation was performed with a precursor isolation window of 1.6 m/z, a maximum injection time of 22 ms, and HCD collision energy of 35%. Monoisotopic-precursor selection was set to “peptide”. Apex detection was not enabled. Precursors were dynamically excluded from resequencing for 20 sec and a mass tolerance of 10 ppm. Precursor ions with charge states that were undetermined, 1, or >5 were excluded. The assay was done with at least 6-replicates pooled together during LC-MS/MS analysis.

### Mass Spectrometry Data Processing

Protein identification and quantification were extracted from raw LC-MS/MS data using the MaxQuant platform v.1.6.3.3 with the Andromeda database searching algorithm and label free quantification (LFQ) algorithm. MaxQuant enables high peptide identification rates, individualized p.p.b.-range mass accuracies and proteome-wide protein quantification (Cox and Mann, 2008, Cox et al., 2014, Tyanova et al., 2016a). Data were searched against a human Uniprot reference database UP0000005640 with 74,468 proteins (August, 2019), a *Porphyromonas gingivalis* database, and a database with other common bacterial species. The false discovery rate (FDR), determined using a reversed database strategy, was set at 1% at the protein and peptide level. Fully tryptic peptides with a minimum of 7 residues were required including cleavage between lysine and proline. Two missed cleavages were permitted. The LFQ feature was on with “Match between runs” enabled for those features that had spectra in at least one of the runs. The “stabilize large ratios” feature was enabled. The first search was performed with a 20 ppm mass tolerance; after recalibration, a 4.5 ppm tolerance was used for the main search. A minimum ratio counts of 2 was required for LFQ protein quantification with at least one unique peptide. Parameters included static modification of cysteine with carbamidomethyl and variable N-terminal acetylation and oxidation of methionine.

The protein groups text file from the MaxQuant search results was processed in Perseus v. 1.6.7.0 (Tyanova et al., 2016b). Identified proteins were filtered to remove proteins only identified by a modified peptide, matches to the reversed database, and potential contaminants. The MQ normalized LFQ intensities for each biological replicate were log2 transformed. Gene ontology annotations were added from a human protein database downloaded from Uniprot with associated GO terms and protein properties of interest.

### Homologous Protein Search and Phylogenetic Tree Construction

All protein signatures identified via Mass Spectrometry were subjected to a NCBI Protein BLAST search using their associated NCBI Protein ID. Each BLAST search was performed under the Position-specific iterated blast algorithm. A total of 20 homologous bacterial proteins ranging from Firmicutes, Bacteroidetes, Actinobacteria and Proteobacteria (including 1 eukaryotic protein if possible) for each protein search were recorded for analysis based on high percent identities (>50%) and low E-value (closest to 0) (Sayers et al., 2022). However, a homologous (significant protein identity) *P. gingivalis* protein could not be identified for *T.* forsythia’s Ragb/SusD protein. Instead *P.gingivalis’* RagB (Protein ID: U2JPV9) was included in the phylogenetic analysis of *T.* forsythia’s Ragb/SusD protein. Phylogenetic analysis of each protein was performed by first constructing multiple sequence alignments using the ClustalW program within the MEGA X application. Each ClustalW alignment was run through the MEGAX Find Best DNA/Protein Model program in MEGA X in order to apply the best evolutionary history model. The Le Gascuel 2008 model was used to construct phylogenetic tree from *P. intermedia’*s and *T. forsythia*’s EF-Tu and *P. intermedia*’s DnaK (Le and Gascuel, 2008). Whelan And Goldman model was used to construct phylogenetic tree from *T. forsythia*’s SusD/RagB protein (Whelan and Goldman, 2001). The maximum likelihood trees were finally generated and exported from 500 bootstrap replicates in the MEGA X application (Tamura et al., 2021, Kumar et al., 2018, Pattengale et al., 2010).

### Functional Regions and Conserved Site Identification

To search for functional conserved sites within homologous proteins, the EXPASY-PROSITE tool and InterPROscan database was utilized (de Castro et al., 2006, Blum et al., 2021). Homologous proteins sequences were uploaded to a CLUSTAL Omega program available through UniProt to generate alignments for annotation (Sievers et al., 2011). Additional known and putative binding substrates for each protein were found from literature review (Sun et al., 2022, Harvey et al., 2019, Tomoyasu et al., 2012, Candela et al., 2010).

### Protein Functional Interaction Prediction

In order to identify known functional interactions within the context of *P. gingivalis*, the STRING database was accessed under confidence mode. *P. gingivalis’* EF-Tu and DnaK were searched within the STRING database. *B. fragilis*’ NanU protein and *T. forsythia’s* RagB/SuSD protein was searched by amino acid sequence within the STRING database. A protein of 98.8% identity (BF1797) was used to construct a STRING network for *T. forsythia’s* RagB/SuSD protein. Only functional interaction networks with a confidence score above 0.7 were displayed and exported (Szklarczyk et al., 2019).

### Assessing the structural homology of select proteins

Published data for the crystallized structures of *P. gingivalis’s* EF-Tu *and T. forsythia’s* RagB/SuSD nutrient uptake outer membrane protein) and thus are no established models available in public depositories. The secondary structure of these proteins was predicted using a predictive structural biology program RosettaTTAFold under Robetta (Yang et al., 2020). Out of the five models generated, the model with the least angstrom error estimates across the protein’s sequence was selected for analysis. Model 3 for *P. gingivalis’* predicted EF-Tu structure and Model 2 for *T. forsythia’s* predicted RagB/SusD family protein was selected from the Robetta provided models. Characterized models for *E.coli*’s EF-Tu (PDB ID:1EFC) and its EF-Tu in complex with *A. baumanni* DsbA (PDB ID: 4P3Y), *P. gingivalis’* RagB (PDB ID: 5CX8) and *B. fragilis* NanU (PBD ID: 4L7T) were acquired from Protein Data Bank (PDB). All models were visualized, annotated and superimposed in UCSF ChimeraX (Goddard et al., 2018, Pettersen et al., 2004, Pettersen et al., 2021).

## 3. Results

### 3.1 Bacterial protein quantities of Elongation Factor Tu (EF-Tu), chaperone protein dnaK and a RagB/SusD increase over a 24 hour P. gingivalis infection

Primary GECs were infected for up to 24 hours with *P. gingivalis* ATCC 33277. After *P. gingivalis* containing vacoules were isolated from infected cells at 3 hour, 6 hour and 24 hour post infection time points, numerous protein signatures belonging to *P. gingivalis, T. forsythia* and *P. intermedia* were identified via mass spectrometry. Label free-quantification (LFQ) was used to measure the amount of specific proteins present from within the vacoules. According to the reported LFQ values of these proteins, the synthesis of homologous intracellular *P. gingivalis* proteins increased over the course of the 24 hour infection (Figure 2). This study will discuss bacterial proteins Elongation Factor Tu (EF-Tu), chaperone protein dnaK and a RagB/SusD nutrient uptake outer membrane protein and their shared properties amongst oral and gut bacteria (Table 1). Mass Spectrometry analysis specifically identified protein signatures residing in Domain 2 of EF-Tu, a nucleotide binding site superfamily domain of dnaK and a SusD-like N terminal domain of the RagB/SusD nutrient uptake outer membrane protein.

**FIGURE 2.**
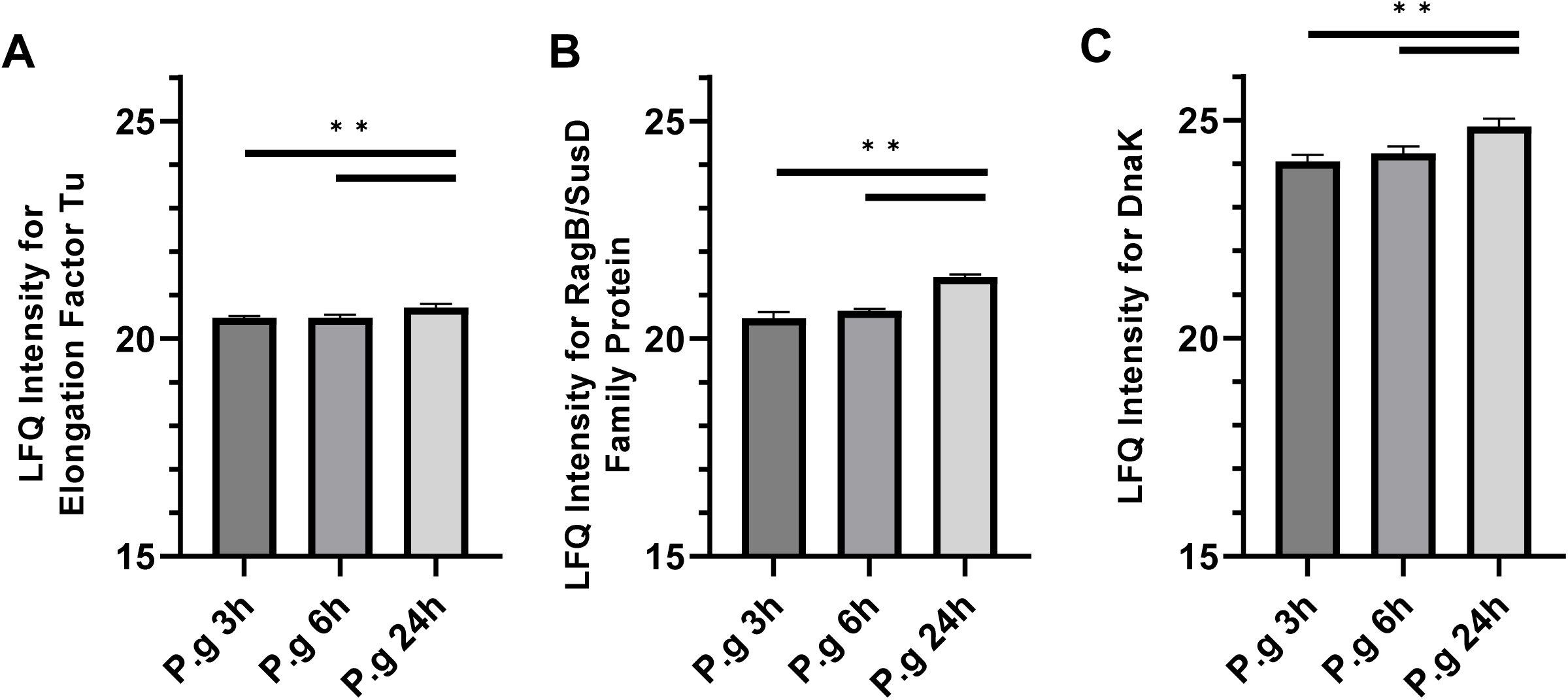
Specific protein signatures for A) Elongation Factor Tu, B) RagB/SusD Family Protein, and C) DnaK, which were originally characterized in other bacterial species and were persistently present in *P. gingivalis-*specific autophagic vacuoles. These autophagic vacuoles were isolated from human primary gingival epithelial cells (GECs) infected with wild-type *P. gingivalis*. GECs were infected with *P. gingivalis* for 3 h, 6 h, or 24 h. The assay was done with at least 6-replicates pooled together during LC-MS/MS analysis.

**TABLE 1.**
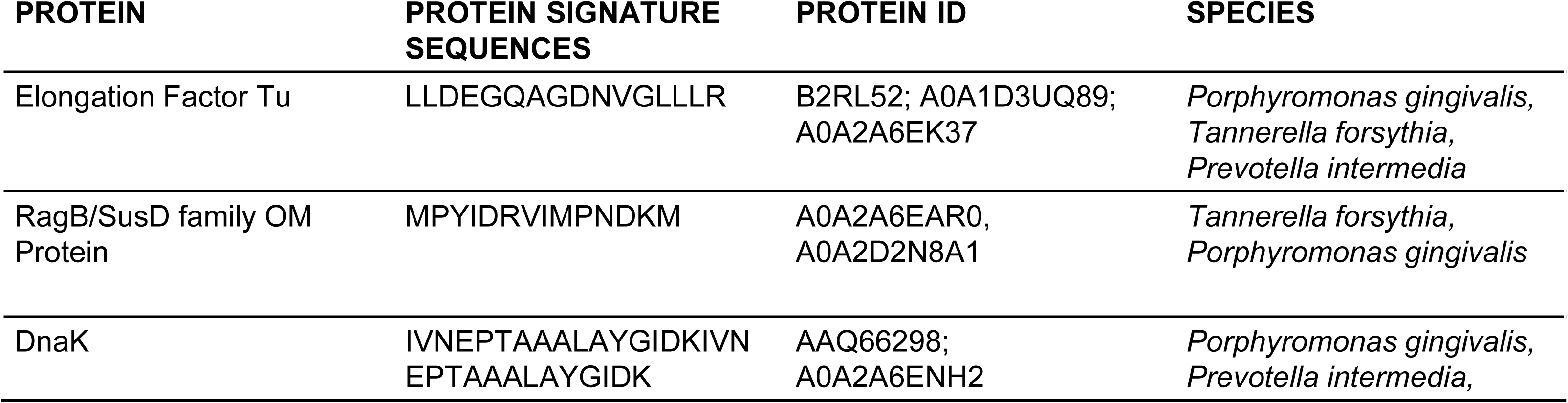
List of protein signatures of interest identified from vacuoles containing *P. gingivalis*. Protein IDs were reported from Mass Spectrometry analysis after identifying specific protein signatures belonging in the biological samples. Homologous *P. gingivalis* proteins containing the protein signatures are displayed as well.

### 3.2 Pathogenic oral bacteria, T. forsythia’s Elongation Factor Tu shares conserved properties with neighboring pathogenic bacteria within the oral microbiome

According to mass spectrometry results, EF-Tu protein signatures belonging to oral bacteria *P. gingivalis*, *T. forsythia*, and *P. intermedia* were identified in the vacuoles isolated from infected primary GEC’s. EF-Tu is a conserved GTPase protein that is intracellularly abundant and essential for protein synthesis across both eukaryotes and prokaryotes (Yikilmaz et al., 2014, Stark et al., 1997). Thus, a number of homologous EF-Tu proteins were identified ranging from bacteria that reside in both oral and gut microbiomes but primarily from bacteria resident in the oral cavity (Supplementary Table 1). In order to assess the extent of relatedness of *T. forsythia’*s EF-Tu with other oral and gut bacteria, phylogenetic analysis was performed. *T. forsythia*’s EF-Tu protein was found to be most closely related to neighboring red oral complex bacteria, *P. gingivalis* from the maximum likelihood tree. Additionally, several homologous EF-Tu proteins from orange oral complex bacteria such as *P. intermedia* and *Fusobacterium nucleatum* had close relation to members of the red oral complex. Red and orange complex bacteria also appear to be extensively related to gut bacteria such as *E. coli, Citrobacter feundii* and *Parabacteriotes distasonis* (Figure 3A). In order to further analyze the relatedness of the *T. forsythia*’s EF-Tu with other bacteria, protein sequences from the same clade were aligned for annotation and SNP detection. A conserved binding site for the ligand GTP was identified via InterPro and denoted in an alignment of these homologous EF-Tu proteins from this closely related gut and oral bacteria (Figure 3A). After extensive literature review, several accessory proteins beyond oral or gut bacterial EF-Tu’s main ligand, GTP, were found to bind to EF-Tu to serve accessory functions (Harvey et al., 2019). These proteins include bacterial DsbA, human holotransferrin, human mucin and human nucleolin (Figure 3B). In summary, these results provide support for the putative and ancestral functional properties of EF-Tu across pathogenic oral bacteria and related bacterial species that reside in different regions of a human host.

**FIGURE 3.**
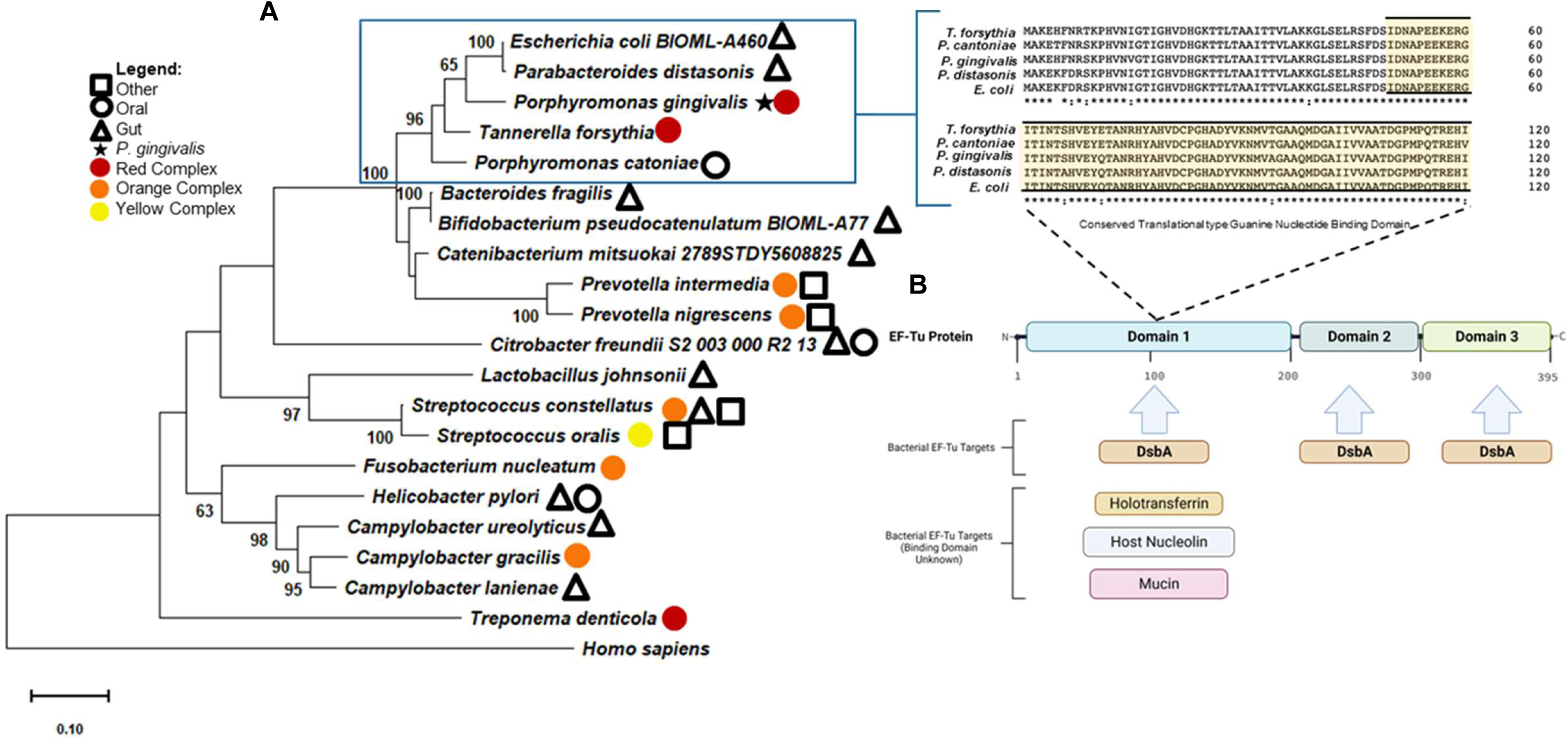
Pathogenic oral complex bacteria share putative accessory binding sites in bacterial EF-TU. (A) Maximum likelihood tree based on *T. forsythia*’s EF-Tu and 20 homologous proteins from oral/gut bacterial species and a control eukaryotic species (n=21). A sequence alignment of *T. forsythia* EF-Tu and closely related species is displayed to the right of the clade. A conserved region shared amongst these species is highlighted in yellow. Bootstrap percentages are hidden if under 50. Red complex bacteria are denoted by a red circle, orange complex bacteria are denoted by an orange circle and yellow complex bacteria are denoted by a yellow circle. Bacteria that can colonize the human gut are denoted by a triangle and bacteria that colonize other parts of the human host are denoted by a square. The black star specifies *P. gingivalis*. (B) Schematic diagram of EF-Tu locus and potential ligands binding to bacterial EF-Tu. GTP is displayed above the sequence alignment binding to the Guanine Nucleotide Binding Domain.

### 3.3 Orange complex bacteria, P. intermedia’s EF-Tu is most related to human gut resident bacteria

A total of 20 homologous proteins were identified via BLAST search ranging from both oral and gut bacteria, most of which were gut resident bacteria. The most homologous proteins to *P. intermedia*’s EF-Tu protein signature were from other orange complex bacteria and bacteria residing in the gut microbiome (e-value closest to 0) (Supplementary Table 2). In concordance with *T. forsythia*’s EF-Tu, EF-Tu in pathogenic oral bacteria are often extensively related to each other (Figure 4). However, in contrast to *T. forsythia*, EF-Tu from *P. intermedia* is primarily related to commensal gut bacteria based on phylogenetic analysis. As in Figure 3A, *P. intermedia’s* and the bacteria that reside within the same clade demonstrate a conserved binding site for GTP indicating a conserved primary function within these closely related bacteria. In conclusion, oral bacteria such as *P. intermedia* may share more proteomic similarities with members of different microbiomes which may serve as evidence for genetic transfer between microbiomes.

**FIGURE 4.**
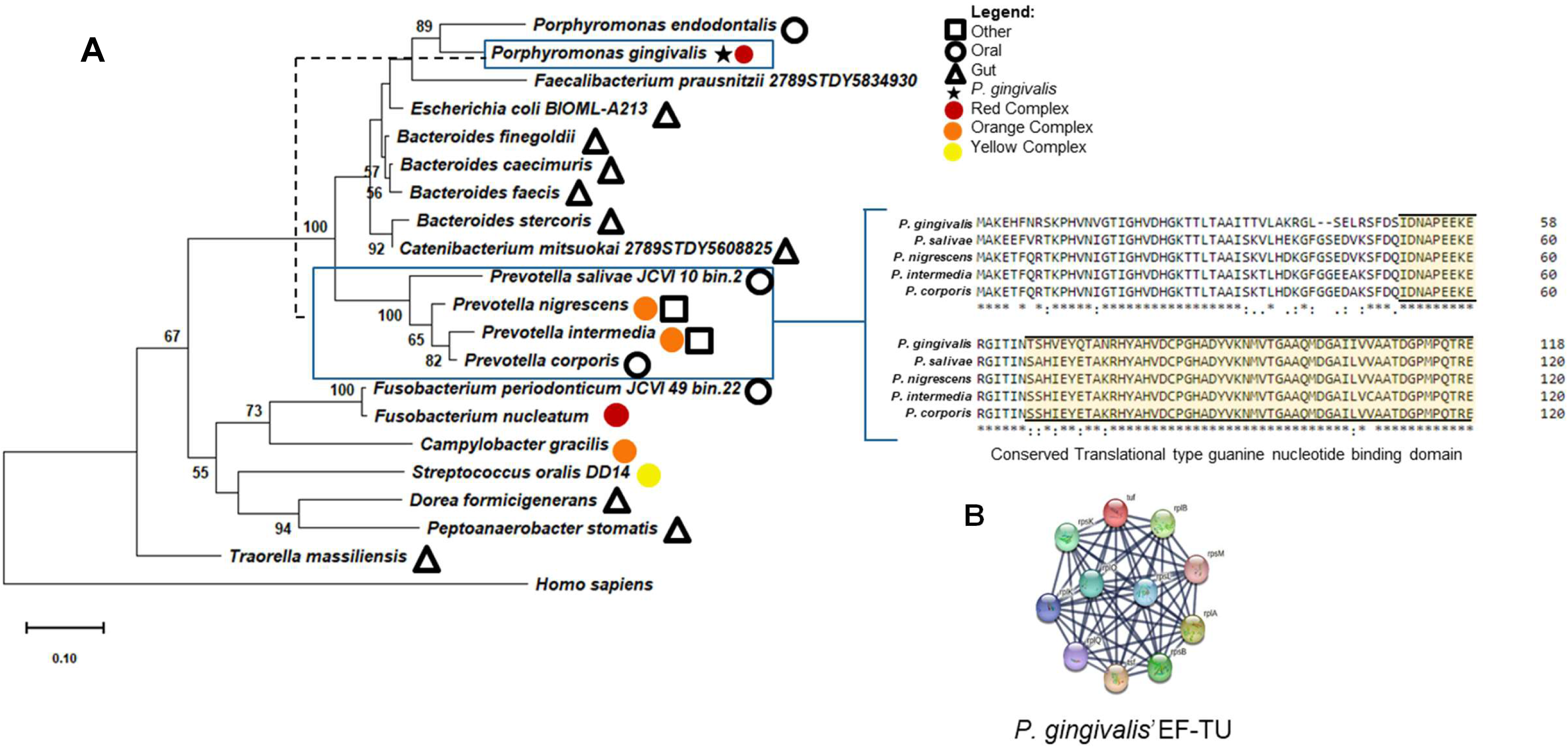
*P. intermedia’s* EF-Tu shares a conserved GTP binding site with gut and oral bacteria. (A) Maximum likelihood tree based on *P. intermedia’s* 20 homologous proteins from oral/gut bacterial species and a control eukaryotic species (n=21). A sequence alignment of *P. intermedia* and closely related species is shown from one clade to the right. A conserved region shared amongst these species is highlighted in yellow. Bootstrap percentages are hidden if under 50. Red complex bacteria are denoted by a red circle, orange complex bacteria are denoted by an orange circle and yellow complex bacteria are denoted by a yellow circle. Bacteria that can colonize the human gut are denoted by a triangle and bacteria that colonize other parts of the human host are denoted by a square. The black star specifies interactions of *P. gingivalis’* EF-Tu. (B) Representative string networks displaying the known functional interactions of *P. gingivalis’* EF-Tu.

### 3.4 P. gingivalis’ EF-Tu structurally homologous to moonlighting EF-Tu from E. coli to moonlighting which may enable pro-survival functions

In order to further assess the relatedness of oral bacterial EF-Tu proteins to members of the gut microbiome, we sought to analysis these proteins structurally. Both *P. gingivalis* and *T.* forsythia’s EF-Tu structure were predicted using a predictive modeling program Robetta to generate possible models with 88% confidence. After these predicted structure were superimposed with *E.coli*’s EF-Tu in its GDP bound state, the two proteins appear to have extensive similarity with a percent homology of 78% (Figure 5A). In addition to there being evidence that these species are related based phylogenetic analysis of these proteins, these proteins shared conserved domains as established in Figure 5B. In a study investigating the biochemical interactions of *Acinetobacter baumannii*’s DsbA enzyme, researchers found that *E. coli*’s EF-Tu can be isolated in complex with *A. baumannii*’s DsbA. To investigate the accessory binding similarities between these two bacteria, we sought to predict if *P. gingivalis’* EF-Tu could potentially bind to *A. baumannii*’s DsbA. After superimposing the predicted structure of *P. gingivalis* with *E.coli*’s EF-Tu bound to *A. baumanni*’s DsbA, it appears that *P. gingivalis* EF-Tu may share the same accessory binding (Figure 5C). To summarize, the moonlighting potential of EF-Tu protein from pathogenic oral bacteria can be predicted via structural homology and this shared moonlighting potential may enable pro-survival functions yet to be studied.

**FIGURE 5.**
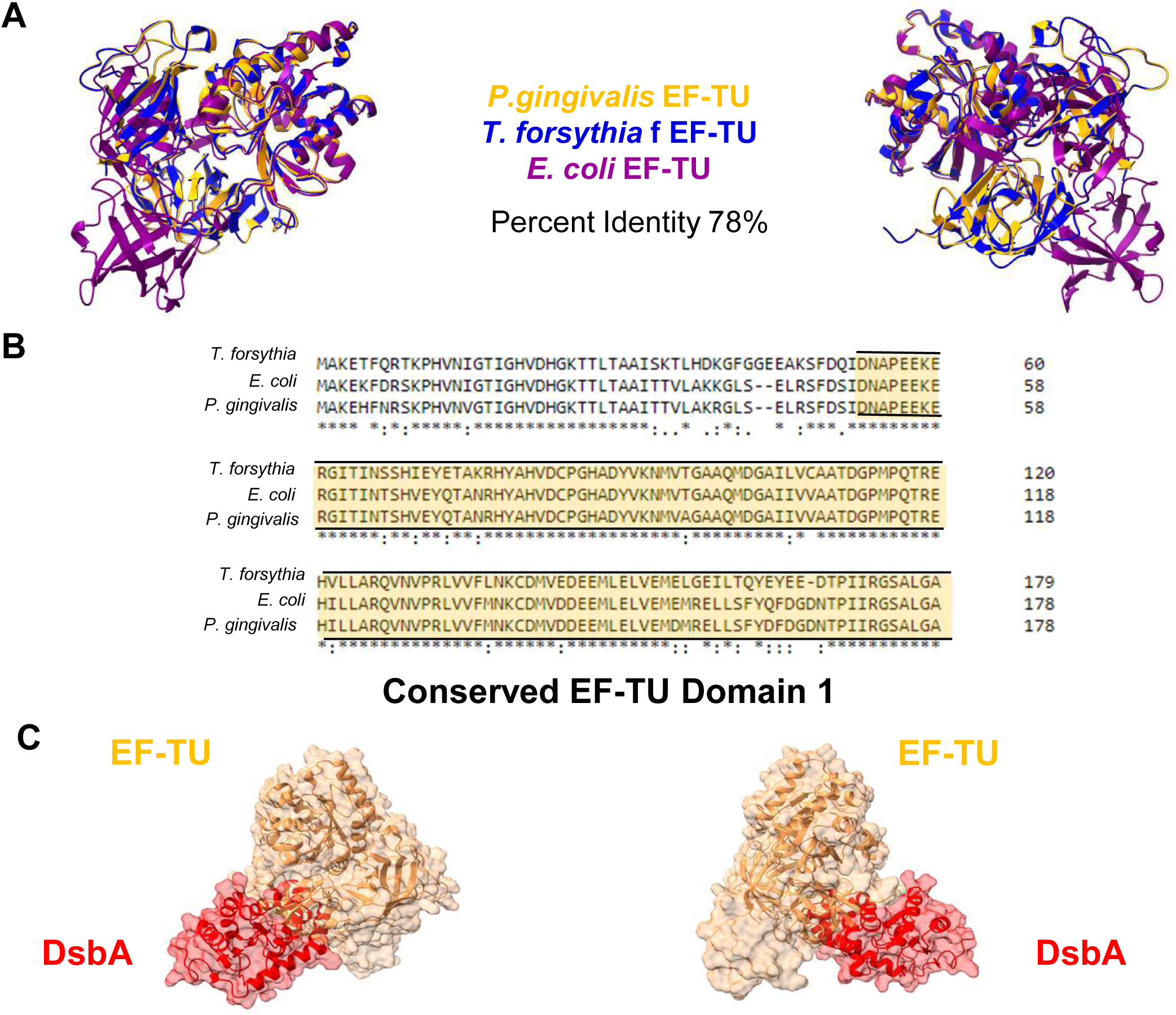
*P. gingivalis’* EF-Tu is structurally similar to *E.coli’s* EF-Tu. (A) Superimposed predicted models of *P. gingivalis’* EF-Tu (Orange Ribbon) with *E.coli’s* EF-TU (Purple Ribbon). (B) A sequence alignment of *P. gingivalis’*, T. forsythia’s and *E. coli’s* EF-Tu protein. A conserved region shared amongst these species is highlighted in yellow. (C) *P. gingivalis’* predicted EFTu (orange ribbon) bound to *A. baumanni’s* DsbA (red ribbon).

### 3.5 T. forsythia’s RagB/SusD family nutrient uptake outer membrane protein is mostly homologous to gut Parabacteriotes

In addition to EF-Tu, a protein signature from the RagB/SusD family was identified from red oral complex bacteria, *T. forsythia*. Rag/SusD proteins are starch binding proteins that allow bacteria to uptake nutrients in their microbial environment and are localized on the outer membrane of *Bacteroidetes* (Glenwright et al., 2017). Phylogenetic analysis revealed that *T. forsythia*’s Rag/SusD protein is most similar to proteins from other *Parabacteriotes* species from the gut. However, this protein was found to be related to other oral bacteria such as orange oral complex bacteria, *P. intermedia* (Supplementary Figure 3, Figure 6). Amongst the closely related proteins, there is a conserved starch binding protein and a lipoprotein lipid attachment domain (Figure 6). This includes a putative haloacid dehalogenase (HAD) protein from P. *gingivalis* which is understood to be involved in metabolism and homeostasis (Tribble et al., 2006). Taken together, this protein derived from oral bacteria, *T. forsythia*, appears to share functional characteristics to gut resident bacteria which may highlight additional conserved or inherited nutrient uptake pathways.

**FIGURE 6.**
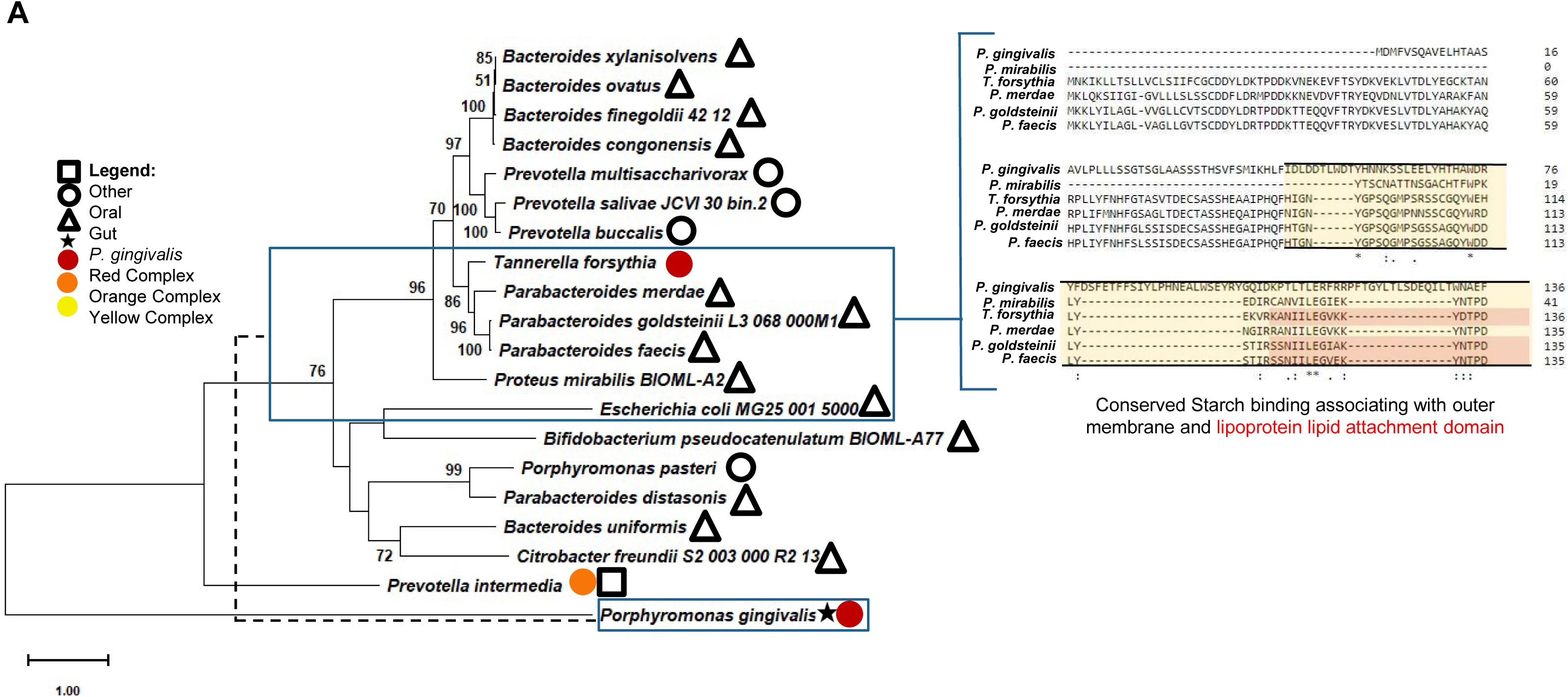
*T. forsythia’s* SusD/RagB like protein share conserved regions with gut bacteria and *P. gingivalis’* RagB. (A) Maximum likelihood tree based on *T. forsythia’s* SusD/RagB protein EF-Tu and 19 homologous proteins from oral/gut bacterial species and a control eukaryotic species (n=20). A sequence alignment of *T. forsythia* and closely related species is shown from one clade is shown to the right. A conserved region shared amongst these species is highlighted in yellow. Bootstrap percentages are hidden if under 50. Red complex bacteria are denoted by a red circle, orange complex bacteria are denoted by an orange circle and yellow complex bacteria are denoted by a yellow circle. Bacteria that can colonize the human gut are denoted by a triangle and bacteria that colonize other parts of the human host are denoted by a square. The black star specifies *P. gingivalis’* RagB protein.

### 3.6 Structural similarities in T. forsythia’s RagB/SusD family protein may enable starch utilization and antigenicity

In order to predict the function of the SusD family protein signature identified from *T, forsythia*, we sought to compare the structures of SusD proteins and RagB protein to the protein of interest. After superimposing *P. gingivalis’* RagB and *T. forsythia’s* RagB/SusD family protein predicted structure with *B. fragilis’* NanU, all three homologous proteins appeared to have similar structures as well. However, despite *P. gingivalis’* RagB protein and *T. forsythia*’s SusD/RagB protein sharing similar structures they exhibit low percent shared identity (Figure 7A-C). Lastly, the predicted structure of RagB/SusD from *T. forsythia* was generated with 72% confidence and appeared to have a wide range of electrostatic potential and a positively charged region outside the putative binding pocket (Figure 7D). After literature review, it was found that the structure of *P. gingivalis’* RagB is extensively similar to SusD of gut bacteria, *Bacteroides thetaiotaomicron* and *Tannerella forsythia’s* NanU (Goulas et al., 2016). In summary, this data indicates that *P. gingivalis’* associated antigen and the nutrient uptake protein exhibits a substantial structural and functional similarity as well as conservation in both oral and gut bacteria.

**FIGURE 7.**
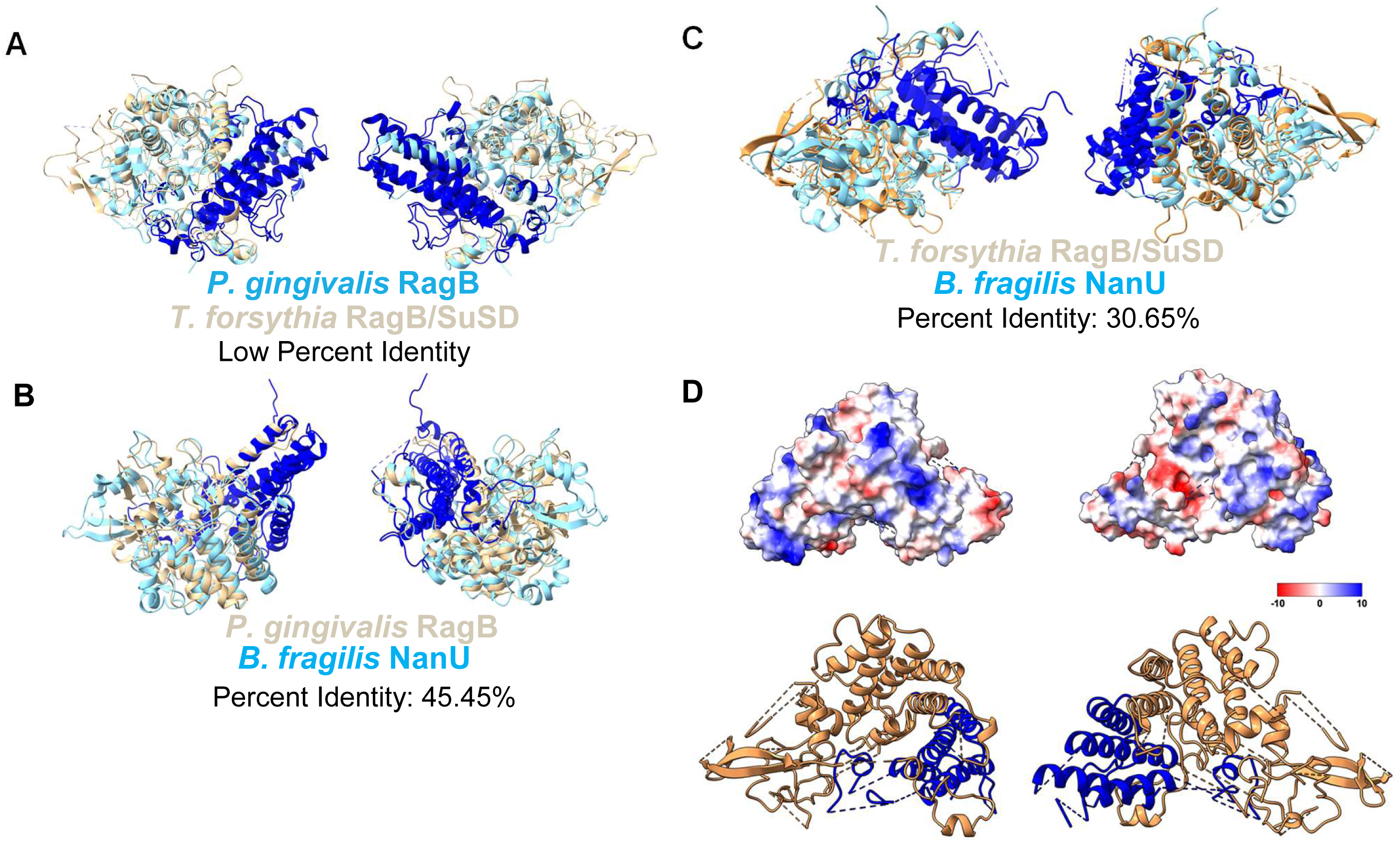
*P. gingivalis’* RagB and *T. forsythia’s* SusD family protein is structurally similar to *Bacillus fragilis’* NanU protein. (A) Superimposed models of *P. gingivalis’* Chain A of RagB (Sky Blue Ribbon) and *T. forsythia’s* predicted Rag/SusD like protein (Beige Ribbon). The starch binding domain is highlighted in dark blue. (B) Superimposed models of *P. gingivalis’* Chain A of RagB (Beige Ribbon) with *B. fragilis* Chain A of NanU (Sky Blue Ribbon). The starch binding domain is highlighted in dark blue. (C) *T. forsythia’s* predicted Rag/SusD like protein (Beige Ribbon) superimposed with *B. fragilis* Chain A of NanU (Sky Blue Ribbon). The conserved starch binding domain is highlighted in dark blue. (D) A ribbon structure and three-dimensional predicted secondary structure image of *T. forsythia’s* EF-Tu (Orange ribbon) depicting the predicted range of surface hydrophobicity. The range in electrostatic potential was calculated using Coulomb’s law. The putative starch binding domain is highlighted in dark blue.

### 3.7 STRING analysis confirms conserved protein functions and predicts putative functions in oral bacteria

In order to determine the known functional interactions of oral bacterial SusD family protein, the STRING database tool was utilized. The known functional interactions of this RagB/SusD protein from *T. forsythia* was found to include connections with a number of putative, hypothetical proteins, a PBS lyase HEAT-like repeat protein and a TonB linked outer membrane protein (Figure 8A). *Bacteroides fragilis*, a gut bacterium that utilizes a SusD like protein called NanU, exhibits similar functional interactions to *T. forsythia*’s RagB/SusD protein (Supplementary Table 6, Figure 8B). In summary, T. *forsythia’s* SusD family protein may enable survival and nutrient uptake functions similar to the gut bacteria, B. fragilis.

**FIGURE 8.**
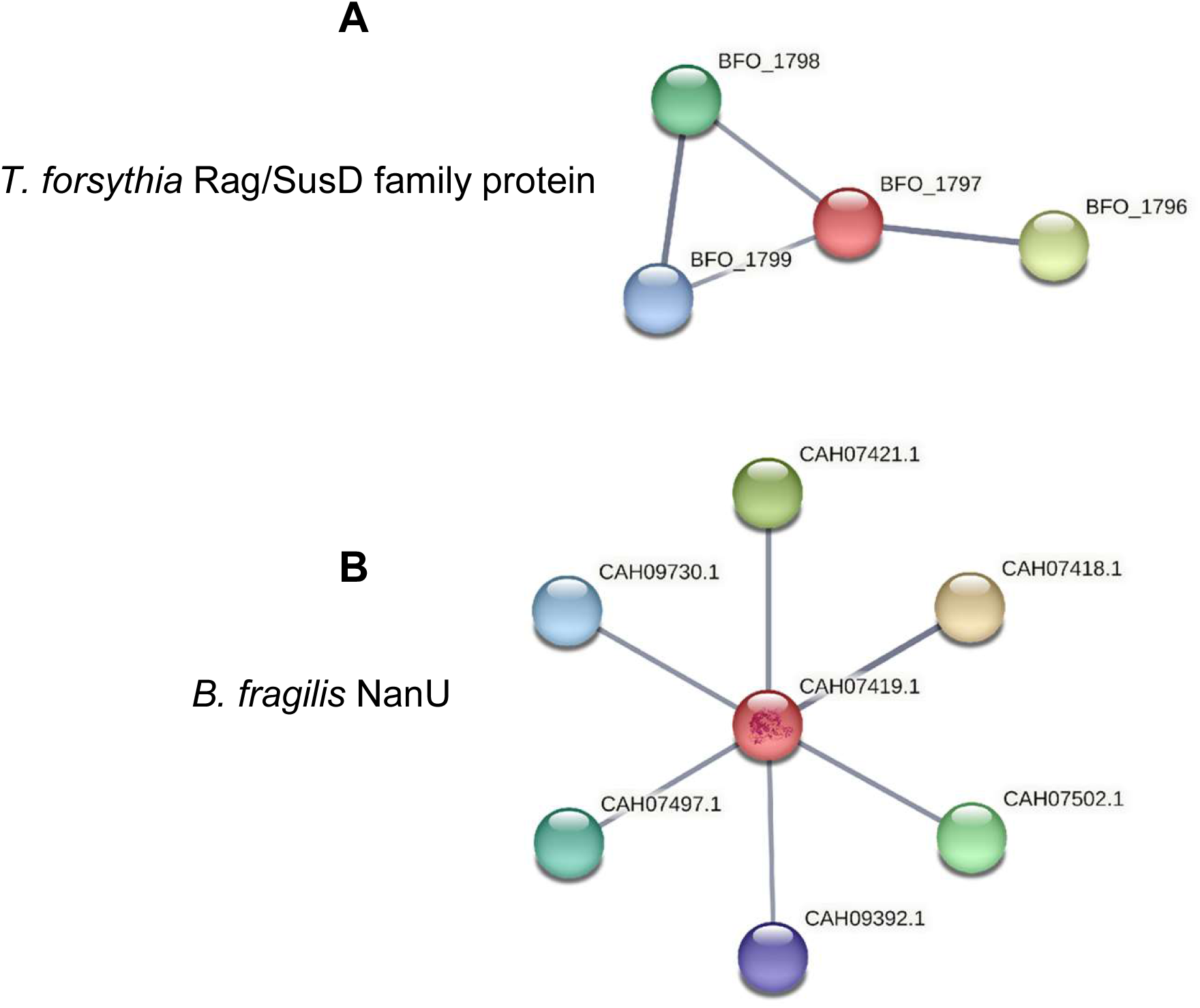
*T. forsythia’s* SusD family protein and *B. fragilis* NanU share similar functional interactions. (A) A string network generated from *T.* forsythia’s Rag/SusD family protein with the Rag/SusD like protein at the center. (B) A string network generated from *B. fragilis’* NanU protein with the NanU protein at the center. The thickness of the “edges” between each node indicates the level of confidence of the interaction between each protein.

### 3.8 Pathogenic Oral bacteria and Gut Bacterial share similar properties in chaperone protein dnaK

A chaperone dnaK protein signature from orange oral complex bacteria *P. intermedia* was also amongst the proteins identified within the *P. gingivalis* contained vacoules. Similar to bacterial EF-Tu, this chaperone protein is essential, and several homologous proteins were identified across oral and gut bacteria (Supplementary Table 4). The most closely related proteins to *P. gingivali*s’ dnaK were from oral bacteria, *P. intermedia* and *P. nigrescens* as well as gut bacteria such as *E. coli*. In addition, red oral complex bacteria *T. denticola* and a number of gut bacteria were shown to have extensive relation to this protein (Figure 9A). Of the homologous proteins most closely related to *P. gingivalis’* dnaK, there are numerous conserved. There are conserved domains shared amongst orange and red complex bacteria such as the conserved Heat Shock protein 70 family signature 1. However, known functional interactions for *P. gingivalis’* dnaK involve other heat shock proteins, chaperone proteins and proteases (Figure 9B). After literature review of chaperone DnaK utility in oral and gut bacteria, it was found that this protein has functional interactions with proteins such as human laminin and plasminogen (Figure 9C). Taken together, these results indicate that oral and gut bacteria have extensive conservation in structure and sequence which may allow for additional pathogenic functions observed in bacteria like *B. lactis* (Candela et al., 2010).

**FIGURE 9.**
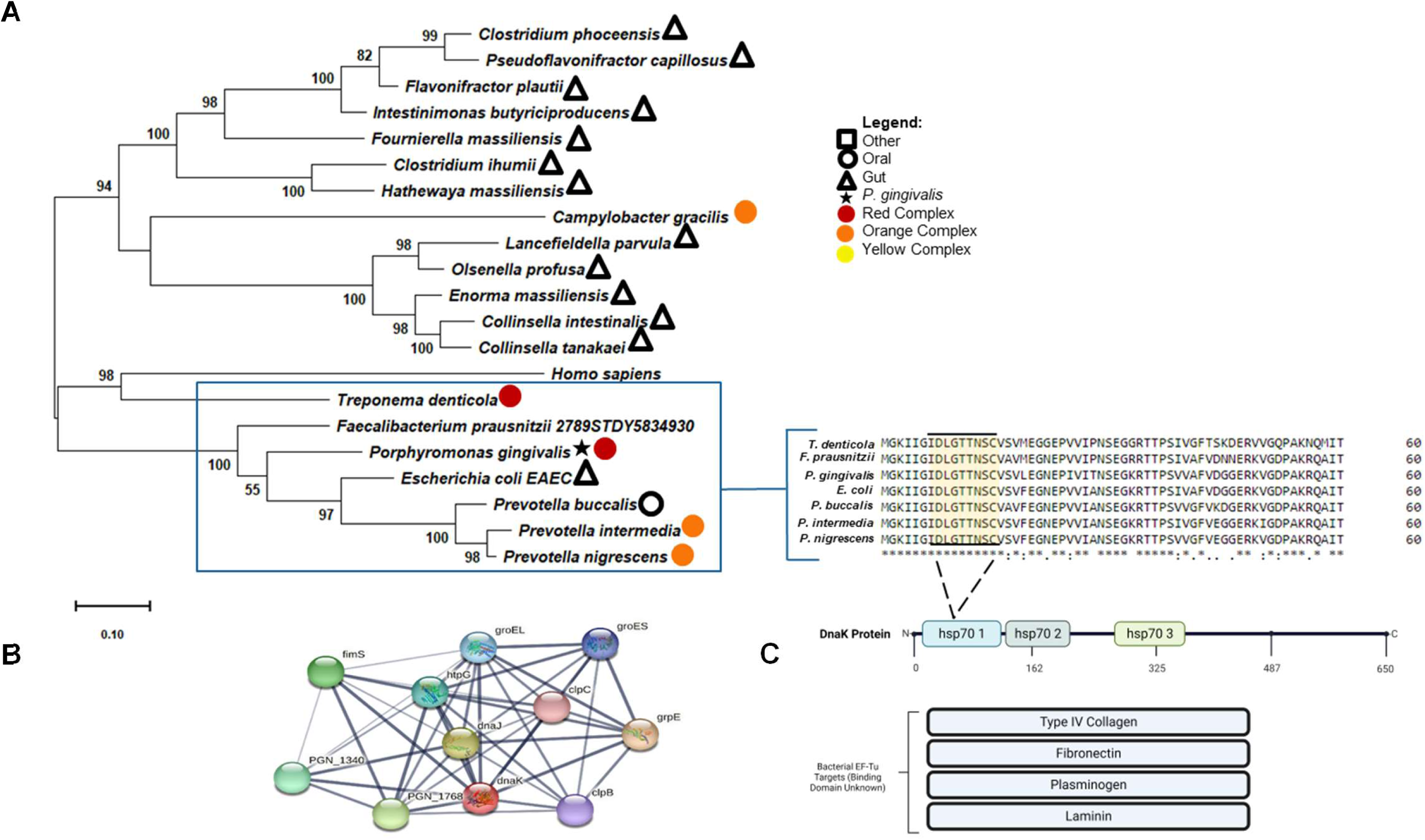
*P. intermedia* DnaK is conserved in pathogenic oral bacteria and gut bacteria. (A) Maximum likelihood tree based on *P. intermedia’s* EF-Tu and 20 homologous proteins from oral/gut bacterial species and a control eukaryotic species (n=21). A sequence alignment of *T. forsythia* and closely related species is shown from one clade is shown to the right. A conserved region shared amongst these species is highlighted in yellow. Bootstrap percentages are hidden if under 50. Red Complex bacteria are denoted in red circle and orange complex are denoted in orange circle. Bacteria that can colonize the human gut are denoted by a triangle and bacteria that colonize other parts of the human host are denoted by a square. The black star specifies *P. gingivalis’*. (B) Schematic diagram of DnaK locus and potential and known ligands. Ligands highlighted in gray indicate there is no literature available for the exact binding on DnaK.

## 4. Discussion

There is little known on the extent of shared survival and pathogenic functions amongst these neighboring bacteria. Understanding the proteomics of one bacterium within the context of one microbiome may aid in our understanding of a related species. Further, assessing the homology of shared proteins may offer potential therapeutic strategies for various clinically relevant oral and gut associated chronic illnesses and pathogens.

Based on phylogenetic analysis of oral bacterial EF-Tu protein signatures identified in *P. gingivalis* contained vacoules, *P. gingivalis’* EF-Tu is closely related to neighboring orange and red oral complex bacteria. As shown in Figure 3A and Figure 4, EF-Tu proteins from oral pathogens, *T. forsythia* or *P. intermedia* are extensively related to pathogenic and commensal oral bacteria. In addition, this study has shown that the evolution of host-adapted bacterial proteomics is not limited to the oral microbiota but extends into the gut bacterial community. For example, EF-Tu proteins from commensal bacteria, *Porphyromonas catoniae*, and commensal gut bacteria, *Bifidobacterium pseudocatenulatum,* appear to be related based on their EF-Tu (Gonzalez-Vazquez et al., 2022). This may suggest genetic transfer between not only oral bacteria but gut resident bacteria which is likely due to the physical connection between the oral cavity and human intestinal tract (Roberts and Kreth, 2014, Guilloux et al., 2021). Moreover, some of these related bacteria can be isolated from not only humans but other animal microbiomes which may suggest potential zoonotic transfer of opportunistic bacteria (Wylensek et al., 2020, Companys et al., 2021, Lagkouvardos et al., 2016) Interestingly, this cytosolic enzyme has enabled similar extracellular virulence and survival functions in various microbiomes of the human host and the potential accessory substrates of bacterial EF-Tu utilized in pathogens are numerous and can be appreciated from Figure 3B (Harvey et al., 2019, Widjaja et al., 2017, Premkumar et al., 2014, Archambaud et al., 2005, Torres et al., 2020). The evolutionary relatedness between bacteria that use EF-Tu as a moonlighting protein and other bacteria may imply that there are uncharacterized extracellular functions for EF-Tu in other oral bacterial pathogens.

In addition to bacterial EF-Tu, the conservation of SusD/RagB outer membrane protein in both oral and gut microbiomes is likely that expression of this protein was transferred between bacteria. In a study examining the evolution of *P. gingivalis*’ RagB protein, they found evidence of long term selection pressure through likelihood ratio tests which is likely as a result of horizontal gene transfer from other Bacteroides bacteria (Su et al., 2010). Our structural analysis as seen in Figure 6B further revealed homology between the RagB/SusD family protein in *T. forsythia* with Chain A of *B. fragilis* NanU protein, a glycan utilization protein. Furthermore, string analysis of *T. forsythia’s* RagB/SusD family proteins and *B. fragilis* NanU protein confirms its utilization in *T. forsythia* as a starch outer membrane binding protein. It was similarly suggested by Hanley et al that the RagAB operon of *P. gingivalis* arose via horizontal gene transfer from a foreign source, potentially from one comprising another component of the periodontal microflora (Hanley et al., 1999). It has also been shown that PgRagB exhibits structural similarity with the *B. thetaiomicron* SusD protein and *T. forsythia* NanU which are both involved in starch and sialic acid uptake (Goulas et al., 2016). Therefore, it is possible that *P. gingivalis* utilizes this conserved starch binding domain in its RagB to acquire sialic acid in a similar manner to that of other species in both the gut and oral microbiomes for later use in several virulence mechanisms. Moreover, *P. gingivalis* utilizes a metabolic protein, HAD phospahatse SerB653, to enable entry into GECs. Given that *T. forsythia*’s SusD/RagB protein is homologous with a putative HAD protein from *P. gingivalis* it possible that these metabolic and nutritional similarities shared amongst oral bacteria may enable virulence and invasiveness (Chen et al., 2001b, Tribble et al., 2006). Overall, oral and gut bacteria appear to utilize similar nutrient uptake systems which further supports a shared evolutionary history.

Furthermore, phylogenetic analysis of DnaK in *P. gingivalis* contained vacoules revealed close relationships to DnaK of other orange complex oral bacteria such as *Prevotella intermedia* and *Prevotella nigrescens*. As previously mentioned, one of the exhibited conserved sites among these homologous proteins was the Heat Shock Protein (HSP) 70 family signature 1. It was hypothesized that the secretion of DnaK may occur due to the ability of *P. intermedia* to escape the oral cavity and colonize in distant parts of the body (Karched et al., 2022), which may be a strong indicator of potential horizontal gene transfer of the DnaK protein between the oral and gut microbiome. Despite possessing several unique characteristics from other known bacterial HSP70s, *P. gingivalis* DnaK was found to behave similarly to *P. intermedia* DnaK in its ability to cooperate with DnaJ and GrpE of the *E. coli* chaperone system (Yoshida et al., 1999). The ability of this protein to cooperate with the DnaJ and GrpE proteins was exhibited further via string network analysis as seen in Figure 8B. Given the homology between these bacteria and the analogous chaperone functioning it is possible that this role and response of DnaK is conserved between species of pathogenic oral and gut bacteria.

Despite there being evidence that oral and gut bacterial protein may share moonlighting functions as a moonlighting factor and mode of adhesion extracellularly, it is important to note that there are limitations to making novel mechanistic conclusions based solely on bioinformatics. Further experimentation is required to conclude whether the binding activities and resulting function of these binding events with EF-Tu can be applied to other bacteria. It is possible that the identification of EF-Tu outside *P. gingivalis* may be an artifact due to upregulated protein synthesis genes while inside the host autophagic vacuoles. In addition, it is understood that *P. gingivalis* can adapt to oral bacterial species communities including *F. nucleatum* and *S. gordonii* by up regulating protein synthesis genes (Kuboniwa et al., 2009). Based on the string network generated in Figure 3B, there is only evidence of *P. gingivalis*’ EF-Tu interacting with ribosomal subunits and the EF-Tu protein which is also required for protein synthesis. The caution must be applied to *P. gingivalis’* RagB protein that is homologous to the SusD family protein shared amongst. Although it may have structural homology with SusD proteins the known function of this protein remains associated with immunogenicity (Potempa et al., 2021).

## 5. Conclusions

Overall, these findings suggest conserved survival and potentially pathogenic mechanisms between oral bacteria, and that this conservation may extend outside the oral microbiome into the gut microbiome. The protein signatures identified within the *P. gingivalis* infected GECs not only characterize the intracellular survival of *P. gingivalis* but also highlight the broad similarities it shares with other pathogenic bacteria. The extent of relatedness between pathogenic oral complex bacteria and pathogenic gut bacteria may provide evidence for the evolution of shared pathogenicity between the two microbiomes. *P. gingivalis* and closely related bacteria use a multitude of virulence mechanisms to enable persistence and invasion within host (Yilmaz, 2008, Lee et al., 2020c, Lee et al., 2018b, Harvey et al., 2019, Yilmaz et al., 2008, Han, 2015, Dashper et al., 2011, Chiu et al., 2017). Further study into the shared survival and virulence pathways amongst oral and gut bacteria may support identification of effective therapeutics and our understanding of established bacteria within microbiomes.

## Funding Source

This work was primarily supported by funding from the NIH grants R01DE030313, F31DE032273, R25GM113278 and T32DE017551.

## Declaration of Competing Interests

Authors declare no conflicts of interest

## Acknowledgments

The authors would like to thank Dr. Lauren Ball, MUSC proteomics core director and MUSC core facilities were supported in part by Hollings Cancer Center Support Grant (P30 CA138313) and the MUSC Digestive Disease Research Cores Center (P30 DK123704).

---

**Supplementary Table 1.**

| ACCESSION # | SPECIES | Percent Identities | Amino Acid # | Region |
| --- | --- | --- | --- | --- |
| WP_014225269.1 | *Tannerella forsythia* |  | 395 | Oral |
| WP_005468460 | *Porphyromonas catoniae* | 92.41 | 395 | Oral |
| WP_121736507.1 | *Parabacteroides distasonis* | 91.86 | 395 | Gut |
| MSK95816.1 | *Escherichia coli* | 91.86 | 395 | Gut |
| WP_195341511 | *Bacteroides fragilis* | 91.14 | 394 | Gut |
| MZM33066 | *Bifidobacterium pseudocatenulatum* | 91.14 | 394 | Gut |
| CUO27731 | *Catenibacterium mitsuokai* | 90.38 | 394 | Gut |
| WP_010955998 | *Porphyromonas gingivalis* | 90.13 | 395 | Oral |
| WP_004366570.1 | *Prevotella nigrescens* | 80.86 | 396 | Oral, urogenital |
| WP_028905370.1 | *Prevotella intermedia* | 80.6 | 396 | Oral, urogenital |
| PZR28947 | *Citrobacter freundii* | 73.86 | 395 | Gut, Oral |
| WP_096015169 | *Campylobacter lanienae* | 71.93 | 399 | Gut |
| WP_001040716 | *Streptococcus oralis* | 71.72 | 398 | Oral, Upper respiratory |
| WP_049480505.1 | *Streptococcus constellatus* | 71.72 | 398 | Oral, Urogenital, Gut |
| WP_024962200 | *Campylobacter ureolyticus* | 71.5 | 399 | Gut |
| WP_001040579.1 | *Helicobacter pylori* | 71.07 | 399 | Oral, Gut |
| WP_032887443.1 | *Fusobacterium nucleatum* | 70.56 | 394 | Oral |
| WP_040304280.1 | *Campylobacter gracilis* | 69.92 | 399 | Oral |
| WP_004894320.1 | *Lactobacillus johnsonii* | 69.47 | 396 | Gut |
| WP_002669993.1 | *Treponema denticola* | 62.72 | 395 | Oral |
| AAB00499 | *Homo sapiens* | 54.68 | 452 | Control |

**Supplementary Table 2.**

| ACCESSION # | SPECIES | Percent Identities | Amino Acid # | Region |
| --- | --- | --- | --- | --- |
| WP_028905370 | *Prevotella intermedia* |  | 396 | Oral, Urogenital |
| WP_025875819 | *Prevotella corporis* | 96.21 | 396 | Oral, Urogenital |
| WP_211791898 | *Prevotella nigrescens* | 93.94 | 396 | Oral |
| MBF1523572 | *Prevotella salivae* | 87.37 | 398 | Oral, Gut |
| WP_007763555 | *Bacteroides finegoldii* | 85.61 | 394 | Gut |
| WP_065539685 | *Bacteroides caecimuris* | 85.1 | 394 | Gut |
| WP_234097660 | *Bacteroides faecis* | 84.6 | 394 | Gut |
| WP_005656566 | *Bacteroides stercoris* | 84.34 | 394 | Gut |
| CUO27731 | *Catenibacterium mitsuokai* | 83.84 | 394 | Gut |
| MZU51045 | *Escherichia coli* | 83.33 | 394 | Gut |
| WP_004335339 | *Porphyromonas endodontalis* | 79.9 | 396 | Oral |
| WP_010955998 | *Porphyromonas gingivalis* | 78.34 | 395 | Oral |
| VDR34100 | *Faecalibacterium prausnitzii* | 76.96 | 395 | Gut |
| MBF1213542 | *Fusobacterium periodonticum* | 70.56 | 395 | Oral |
| WP_032887443 | *Fusobacterium nucleatum* | 70.05 | 394 | Oral |
| WP_118358359 | *Dorea formicigenerans* | 69.6 | 397 | Gut |
| WP_071441959 | *Traorella massiliensis* | 69.54 | 394 | Gut |
| WP_009527606 | *Peptoanaerobacter stomatis* | 68.84 | 397 | Oral |
| WP_040304280 | *Campylobacter gracilis* | 67.92 | 399 | Oral |
| KXT83818 | *Streptococcus oralis* | 66.92 | 398 | Oral |
| AAB00499 | *Homo sapiens* | 54.20 | 452 | Control |

**Supplementary Table 3.**

| ACCESSION # | SPECIES | Percent Identities | Amino Acid # | Region |
| --- | --- | --- | --- | --- |
| WP_097530727 | *Tannerella forsythia* |  | 631 | Oral |
| WP_106831261 | *Parabacteroides pacaensis* | 74.96 | 630 | Gut |
| MBS6574981 | *Parabacteroides goldsteinii* | 69.15 | 627 | Gut |
| WP_183668678 | *Parabacteroides faecis* | 68.76 | 627 | Gut |
| WP_195292649 | *Parabacteroides merdae* | 67.19 | 634 | Gut |
| GAP81744.1 | *Porphyromonas gingivalis* | 64.29 | 503 | Oral |
| WP_007573186 | *Prevotella multisaccharivorax* | 52.66 | 645 | Oral |
| WP_004351005 | *Prevotella buccalis* | 52.45 | 642 | Oral |
| MBF1544323 | *Prevotella salivae* | 52.14 | 642 | Oral |
| WP_240062496 | *Bacteroides xylanisolvens* | 52.02 | 636 | Gut |
| OKZ23327 | *Bacteroides finegoldii* | 51.93 | 642 | Gut |
| WP_022136618 | *Bacteroides congonensis* | 51.91 | 642 | Gut |
| WP_074637268 | *Bacteroides ovatus* | 51.85 | 642 | Gut |
| MTT48421 | *Proteus mirabilis* | 43.88 | 370 | Gut |
| MZM33202 | *Bifidobacterium pseudocatenulatum* | 29.41 | 240 | Gut |
| WP_195207990 | *Bacteroides uniformis* | 28.73 | 563 | Gut |
| PZR29998 | *Citrobacter freundii* | 27.00 | 579 | Gut |
| WP_135958631 | *Parabacteroides distasonis* | 24.88 | 535 | Gut |
| PFX22450 | *Stylophora pistillata* | 24.66 | 3185 | Control |
| MBL1007498 | *Escherichia coli* | 24.5 | 654 | Gut |
| WP_188807090 | *Porphyromonas pasteri* | 23.6 | 548 | Oral |
| WP_028906323 | *Prevotella intermedia* | 22.39 | 544 | Oral |

**Supplementary Table 4.**
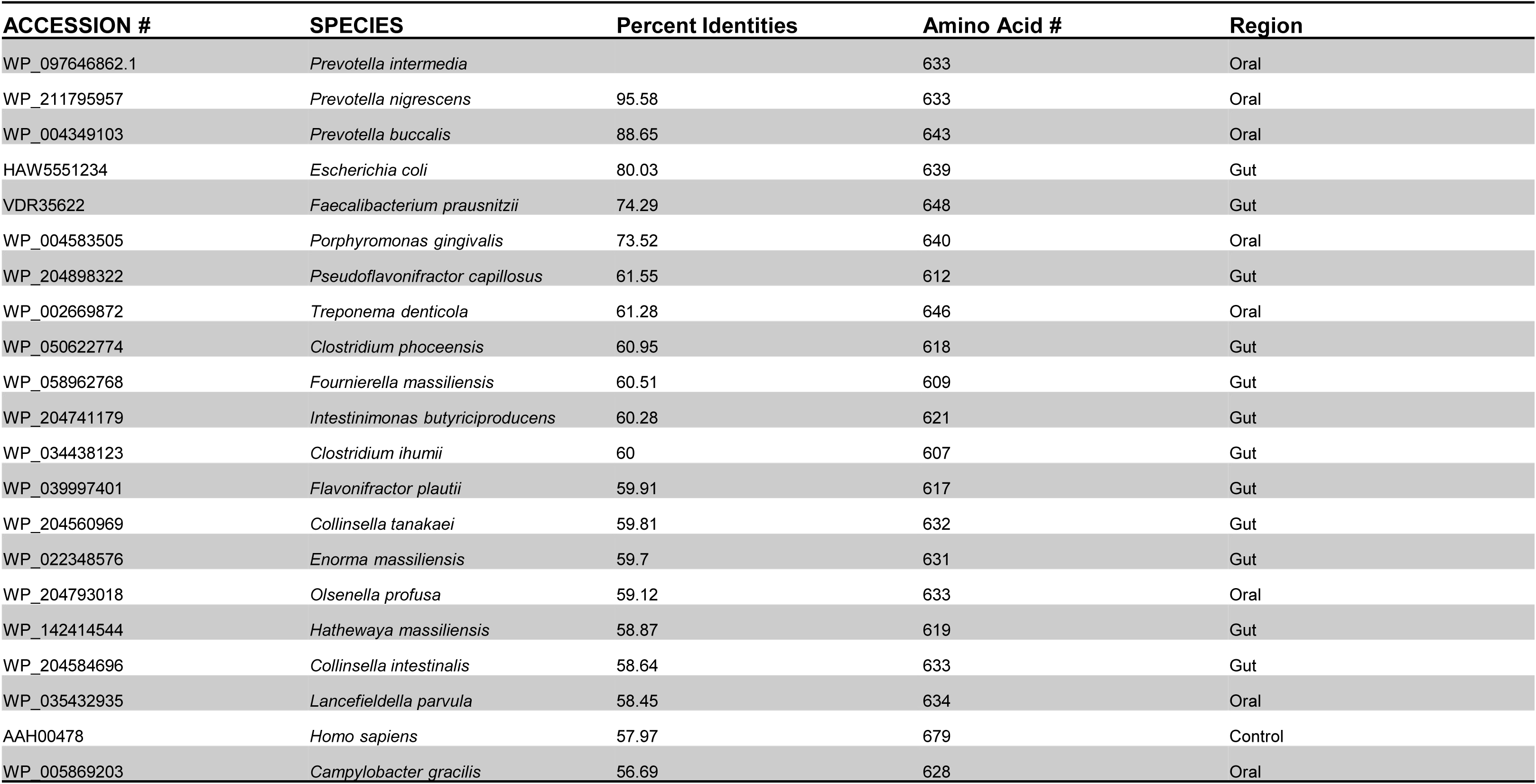

**Supplementary Table 5.**

| Node | Protein | Confidence Score |
| --- | --- | --- |
| CAH07419.1 | NanU Protein | 0.706-0.929 |
| CAH07421.1 | Similar to Bacteroides fragilis outer membrane protein omp121 | 0.769 |
| CAH07497.1 | Putative outer membrane protein | 0.764 |
| CAH07502.1 | Starch-binding outer membrane protein | 0.765 |
| CAH09392.1 | Putative export protein | 0.706 |
| CAH09730.1 | Putative outer membrane protein | 0.756 |

**Supplementary Table 6.**

| **Node** | **Protein** | **Confidence Score** |
| --- | --- | --- |
| BFO_1797 | Rag/SusD Family Protein (Vesicle Isolation) | 0.707-0.912 |
| BFO_1796 | TonB linked outer membrane protein | 0.912 |
| BFO_1798 | Tat pathway signal sequence domain protein | 0.734-0.925 |
| BFO_1799 | Putative lipoprotein | 0.707-0.925 |

## Notes

### Competing Interest Statement

The authors have declared no competing interest.

### Summary of Updates

Minor edits were made to the abstract for clarification.

## References

Archambaud, C., Gouin, E., Pizarro-Cerda, J., Cossart, P. & Dussurget, O. 2005. Translation elongation factor EF-Tu is a target for Stp, a serine-threonine phosphatase involved in virulence of Listeria monocytogenes. Mol Microbiol, 56, 383–96.

Atanasova, K., Lee, J., Roberts, J., Lee, K., Ojcius, D. M. & Yilmaz, Ö. 2016. Nucleoside-Diphosphate-Kinase of P. gingivalis is Secreted from Epithelial Cells In the Absence of a Leader Sequence Through a Pannexin-1 Interactome. Scientific Reports, 6, 37643.

Atanasova, K. R. & Yilmaz, Ö. 2015. Prelude to oral microbes and chronic diseases: past, present and future. Microbes and Infection, 17, 473–483.

Avila, M., Ojcius, D. M. & Yilmaz, O. 2009. The oral microbiota: living with a permanent guest. DNA Cell Biol, 28, 405–11.

Blum, M., Chang, H. Y., Chuguransky, S., Grego, T., Kandasaamy, S., Mitchell, A., Nuka, G., Paysan-Lafosse, T., Qureshi, M., Raj, S., Richardson, L., Salazar, G. A., Williams, L., Bork, P., Bridge, A., Gough, J., Haft, D. H., Letunic, I., Marchler-Bauer, A., Mi, H., Natale, D. A., Necci, M., Orengo, C. A., Pandurangan, A. P., Rivoire, C., Sigrist, C. J. A., Sillitoe, I., Thanki, N., Thomas, P. D., Tosatto, S. C. E., Wu, C. H., Bateman, A. & Finn, R. D. 2021. The InterPro protein families and domains database: 20 years on. Nucleic Acids Res, 49, D344–D354.

Candela, M., Centanni, M., Fiori, J., Biagi, E., Turroni, S., Orrico, C., Bergmann, S., Hammerschmidt, S. & Brigidi, P. 2010. DnaK from Bifidobacterium animalis subsp. lactis is a surface-exposed human plasminogen receptor upregulated in response to bile salts. Microbiology (Reading), 156, 1609–1618.

Chen, T., Nakayama, K., Belliveau, L. & Duncan, M. J. 2001a. Porphyromonas gingivalis gingipains and adhesion to epithelial cells. Infection and immunity, 69, 3048–3056.

Chen, W., Laidig, K. E., Park, Y., Park, K., Yates, J. R., 3rd, Lamont, R. J. & Hackett, M. 2001b. Searching the Porphyromonas gingivalis genome with peptide fragmentation mass spectra. Analyst, 126, 52–7.

Chiu, K.-H., Wang, L.-H., Tsai, T.-T., Lei, H.-Y. & Liao, P.-C. 2017. Secretomic Analysis of Host–Pathogen Interactions Reveals That Elongation Factor-Tu Is a Potential Adherence Factor of Helicobacter pylori during Pathogenesis. Journal of Proteome Research, 16, 264–273.

Chowdhury, N., Wellslager, B., Lee, H., Gilbert, J. L. & Yilmaz, Ö. 2024. Glutamate is a key nutrient for Porphyromonas gingivalis growth and survival during intracellular autophagic life under nutritionally limited conditions. bioRxiv, 2024.07.08.602514.

Companys, J., Gosalbes, M. J., Pla-Paga, L., Calderon-Perez, L., Llaurado, E., Pedret, A., Valls, R. M., Jimenez-Hernandez, N., Sandoval-Ramirez, B. A., Del Bas, J. M., Caimari, A., Rubio, L. & Sola, R. 2021. Gut Microbiota Profile and Its Association with Clinical Variables and Dietary Intake in Overweight/Obese and Lean Subjects: A Cross-Sectional Study. Nutrients, 13.

Cox, J., Hein, M. Y., Luber, C. A., Paron, I., Nagaraj, N. & Mann, M. 2014. Accurate proteome-wide label-free quantification by delayed normalization and maximal peptide ratio extraction, termed MaxLFQ. Mol Cell Proteomics, 13, 2513–26.

Cox, J. & Mann, M. 2008. MaxQuant enables high peptide identification rates, individualized p.p.b.-range mass accuracies and proteome-wide protein quantification. Nat Biotechnol, 26, 1367–72.

Dashper, S. G., Seers, C. A., Tan, K. H. & Reynolds, E. C. 2011. Virulence factors of the oral spirochete Treponema denticola. Journal of dental research, 90, 691–703.

de Castro, E., Sigrist, C. J., Gattiker, A., Bulliard, V., Langendijk-Genevaux, P. S., Gasteiger, E., Bairoch, A. & Hulo, N. 2006. ScanProsite: detection of PROSITE signature matches and ProRule-associated functional and structural residues in proteins. Nucleic Acids Res, 34, W362–5.

Dewhirst, F. E., Chen, T., Izard, J., Paster, B. J., Tanner, A. C., Yu, W. H., Lakshmanan, A. & Wade, W. G. 2010. The human oral microbiome. J Bacteriol, 192, 5002–17.

Faran Ali, S. M. & Tanwir, F. 2012. Oral microbial habitat a dynamic entity. Journal of Oral Biology and Craniofacial Research, 2, 181–187.

Glenwright, A. J., Pothula, K. R., Bhamidimarri, S. P., Chorev, D. S., Basle, A., Firbank, S. J., Zheng, H., Robinson, C. V., Winterhalter, M., Kleinekathofer, U., Bolam, D. N. & Van Den Berg, B. 2017. Structural basis for nutrient acquisition by dominant members of the human gut microbiota. Nature, 541, 407–411.

Goddard, T. D., Huang, C. C., Meng, E. C., Pettersen, E. F., Couch, G. S., Morris, J. H. & Ferrin, T. E. 2018. UCSF ChimeraX: Meeting modern challenges in visualization and analysis. Protein Sci, 27, 14–25.

Gonzalez-Vazquez, R., Zuniga-Leon, E., Torres-Maravilla, E., Leyte-Lugo, M., Mendoza-Perez, F., Hernandez-Delgado, N. C., Perez-Pasten-Borja, R., Azaola-Espinosa, A. & Mayorga-Reyes, L. 2022. Genomic and Biochemical Characterization of Bifidobacterium pseudocatenulatum JCLA3 Isolated from Human Intestine. Microorganisms, 10.

Goulas, T., Garcia-Ferrer, I., Hutcherson, J. A., Potempa, B. A., Potempa, J., Scott, D. A. & Gomis-Ruth, F. X. 2016. Structure of RagB, a major immunodominant outer-membrane surface receptor antigen of Porphyromonas gingivalis. Mol Oral Microbiol, 31, 472–485.

Guilloux, C. A., Lamoureux, C., Beauruelle, C. & Hery-Arnaud, G. 2021. Porphyromonas: A neglected potential key genus in human microbiomes. Anaerobe, 68, 102230.

Haffajee, A. D. & Socransky, S. S. 1994. Microbial etiological agents of destructive periodontal diseases. Periodontol 2000, 5, 78–111.

Haffajee, A. D. & Socransky, S. S. 2005. Microbiology of periodontal diseases: introduction. Periodontol 2000, 38, 9–12.

Hajishengallis, G., Darveau, R. P. & Curtis, M. A. 2012. The keystone-pathogen hypothesis. Nat Rev Microbiol, 10, 717–25.

Han, Y. W. 2015. Fusobacterium nucleatum: a commensal-turned pathogen. Current opinion in microbiology, 23, 141–147.

Hanley, S. A., Aduse-Opoku, J. & Curtis, M. A. 1999. A 55-kilodalton immunodominant antigen of Porphyromonas gingivalis W50 has arisen via horizontal gene transfer. Infect Immun, 67, 1157–71.

Harvey, K. L., Jarocki, V. M., Charles, I. G. & Djordjevic, S. P. 2019. The Diverse Functional Roles of Elongation Factor Tu (EF-Tu) in Microbial Pathogenesis. Front Microbiol, 10, 2351.

Human Microbiome Project, C. 2012. Structure, function and diversity of the healthy human microbiome. Nature, 486, 207–214.

Jenkinson, H. F. & Lamont, R. J. 2005. Oral microbial communities in sickness and in health. Trends Microbiol, 13, 589–95.

Jia, L., Han, N., Du, J., Guo, L., Luo, Z. & Liu, Y. 2019. Pathogenesis of Important Virulence Factors of Porphyromonas gingivalis via Toll-Like Receptors. Frontiers in cellular and infection microbiology, 9, 262–262.

Jiao, Y., Hasegawa, M. & Inohara, N. 2014. The Role of Oral Pathobionts in Dysbiosis during Periodontitis Development. Journal of Dental Research, 93, 539–546.

Karched, M., Bhardwaj, R. G., Qudeimat, M., Al-Khabbaz, A. & Ellepola, A. 2022. Proteomic analysis of the periodontal pathogen Prevotella intermedia secretomes in biofilm and planktonic lifestyles. Sci Rep, 12, 5636.

Kassebaum, N. J., Smith, A. G. C., Bernabé, E., Fleming, T. D., Reynolds, A. E., Vos, T., Murray, C. J. L. & Marcenes, W. 2017. Global, Regional, and National Prevalence, Incidence, and Disability-Adjusted Life Years for Oral Conditions for 195 Countries, 1990–2015: A Systematic Analysis for the Global Burden of Diseases, Injuries, and Risk Factors. Journal of Dental Research, 96, 380–387.

Kuboniwa, M., Hendrickson, E. L., Xia, Q., Wang, T., Xie, H., Hackett, M. & Lamont, R. J. 2009. Proteomics of Porphyromonas gingivalis within a model oral microbial community. BMC Microbiol, 9, 98.

Kumar, S., Stecher, G., Li, M., Knyaz, C. & Tamura, K. 2018. MEGA X: Molecular Evolutionary Genetics Analysis across Computing Platforms. Mol Biol Evol, 35, 1547–1549.

Lagkouvardos, I., Pukall, R., Abt, B., Foesel, B. U., Meier-Kolthoff, J. P., Kumar, N., Bresciani, A., Martinez, I., Just, S., Ziegler, C., Brugiroux, S., Garzetti, D., Wenning, M., Bui, T. P., Wang, J., Hugenholtz, F., Plugge, C. M., Peterson, D. A., Hornef, M. W., Baines, J. F., Smidt, H., Walter, J., Kristiansen, K., Nielsen, H. B., Haller, D., Overmann, J., Stecher, B. & Clavel, T. 2016. The Mouse Intestinal Bacterial Collection (miBC) provides host-specific insight into cultured diversity and functional potential of the gut microbiota. Nat Microbiol, 1, 16131.

Le, S. Q. & Gascuel, O. 2008. An improved general amino acid replacement matrix. Mol Biol Evol, 25, 1307–20.

Lee, J., Roberts, J. S., Atanasova, K. R., Chowdhury, N. & Yilmaz, Ö. 2018a. A novel kinase function of a nucleoside-diphosphate-kinase homologue in Porphyromonas gingivalis is critical in subversion of host cell apoptosis by targeting heat-shock protein 27. Cell Microbiol, 20, e12825.

Lee, J., Spooner, R., Chowdhury, N., Pandey, V., Wellslager, B., Atanasova, K., Evans, Z. & Yilmaz, O. 2020a. In Situ Intraepithelial Localizations of Opportunistic Pathogens, Porphyromonas gingivalis and Filifactor alocis, in Human Gingiva. Current Research in Microbial Sciences. Current Research in Microbial Sciences, 1, 7–17.

Lee, J. S., Chowdhury, N., Roberts, J. S. & Yilmaz, O. 2020b. Host surface ectonucleotidase-CD73 and the opportunistic pathogen, Porphyromonas gingivalis, cross-modulation underlies a new homeostatic mechanism for chronic bacterial survival in human epithelial cells. Virulence, 11, 414–429.

Lee, J. S., Chowdhury, N., Roberts, J. S. & Yilmaz, Ö. 2020c. Host surface ectonucleotidase-CD73 and the opportunistic pathogen, Porphyromonas gingivalis, cross-modulation underlies a new homeostatic mechanism for chronic bacterial survival in human epithelial cells. Virulence, 11, 414–429.

Lee, J. S., Spooner, R., Chowdhury, N., Pandey, V., Wellslager, B., Atanasova, K. R., Evans, Z. & Yilmaz, Ö. 2020d. In Situ Intraepithelial Localizations of Opportunistic Pathogens, Porphyromonas gingivalis and Filifactor alocis, in Human Gingiva. Current Research in Microbial Sciences, 1, 7–17.

Lee, J. S. & Yilmaz, Ö. 2020. Key Elements of Gingival Epithelial Homeostasis upon Bacterial Interaction. Journal of Dental Research, 100, 333–340.

Lee, K., Roberts, J. S., Choi, C. H., Atanasova, K. R. & Yilmaz, O. 2018b. Porphyromonas gingivalis traffics into endoplasmic reticulum-rich-autophagosomes for successful survival in human gingival epithelial cells. Virulence, 9, 845–859.

Lee, Y.-H., Chung, S. W., Auh, Q. S., Hong, S.-J., Lee, Y.-A., Jung, J., Lee, G.-J., Park, H. J., Shin, S.-I. & Hong, J.-Y. 2021. Progress in Oral Microbiome Related to Oral and Systemic Diseases: An Update. Diagnostics (Basel, Switzerland), 11, 1283.

Mohanty, R., Asopa, S. J., Joseph, M. D., Singh, B., Rajguru, J. P., Saidath, K. & Sharma, U. 2019. Red complex: Polymicrobial conglomerate in oral flora: A review. Journal of family medicine and primary care, 8, 3480–3486.

Nakayama, M. & Ohara, N. 2017. Molecular mechanisms of Porphyromonas gingivalis-host cell interaction on periodontal diseases. The Japanese dental science review, 53, 134–140.

Nakhjiri, S. F., Park, Y., Yilmaz, O., Chung, W. O., Watanabe, K., El-Sabaeny, A., Park, K. & Lamont, R. J. 2001. Inhibition of epithelial cell apoptosis by Porphyromonas gingivalis. FEMS Microbiol Lett, 200, 145–9.

Narengaowa, Kong, W., Lan, F., Awan, U. F., Qing, H. & Ni, J. 2021. The Oral-Gut-Brain AXIS: The Influence of Microbes in Alzheimer’s Disease. Frontiers in Cellular Neuroscience, 15.

Pattengale, N. D., Alipour, M., Bininda-Emonds, O. R., Moret, B. M. & Stamatakis, A. 2010. How many bootstrap replicates are necessary? J Comput Biol, 17, 337–54.

Pettersen, E. F., Goddard, T. D., Huang, C. C., Couch, G. S., Greenblatt, D. M., Meng, E. C. & Ferrin, T. E. 2004. UCSF Chimera--a visualization system for exploratory research and analysis. J Comput Chem, 25, 1605–12.

Pettersen, E. F., Goddard, T. D., Huang, C. C., Meng, E. C., Couch, G. S., Croll, T. I., Morris, J. H. & Ferrin, T. E. 2021. UCSF ChimeraX: Structure visualization for researchers, educators, and developers. Protein Sci, 30, 70–82.

Potempa, J., Madej, M. & Scott, D. A. 2021. The RagA and RagB proteins of Porphyromonas gingivalis. Mol Oral Microbiol, 36, 225–232.

Premkumar, L., Kurth, F., Duprez, W., Groftehauge, M. K., King, G. J., Halili, M. A., Heras, B. & Martin, J. L. 2014. Structure of the Acinetobacter baumannii dithiol oxidase DsbA bound to elongation factor EF-Tu reveals a novel protein interaction site. J Biol Chem, 289, 19869–80.

Roberts, A. P. & Kreth, J. 2014. The impact of horizontal gene transfer on the adaptive ability of the human oral microbiome. Front Cell Infect Microbiol, 4, 124.

Roberts, J. S., Atanasova, K. R., Lee, J., Diamond, G., Deguzman, J., Hee Choi, C. & Yilmaz, O. 2017a. Opportunistic Pathogen Porphyromonas gingivalis Modulates Danger Signal ATP-Mediated Antibacterial NOX2 Pathways in Primary Epithelial Cells. Front Cell Infect Microbiol, 7, 291.

Roberts, J. S., Atanasova, K. R., Lee, J., Diamond, G., Deguzman, J., Hee Choi, C. & Yilmaz, Ö. 2017b. Opportunistic Pathogen Porphyromonas gingivalis Modulates Danger Signal ATP-Mediated Antibacterial NOX2 Pathways in Primary Epithelial Cells. Frontiers in Cellular and Infection Microbiology, 7.

Sayers, E. W., Bolton, E. E., Brister, J. R., Canese, K., Chan, J., Comeau, D. C., Connor, R., Funk, K., Kelly, C., Kim, S., Madej, T., Marchler-Bauer, A., Lanczycki, C., Lathrop, S., Lu, Z., Thibaud-Nissen, F., Murphy, T., Phan, L., Skripchenko, Y., Tse, T., Wang, J., Williams, R., Trawick, B. W., Pruitt, K. D. & Sherry, S. T. 2022. Database resources of the national center for biotechnology information. Nucleic Acids Res, 50, D20–D26.

Shaw, L. P., Smith, A. M. & Roberts, A. P. 2017. The oral microbiome. Emerging Topics in Life Sciences, 1, 287–296.

Sievers, F., Wilm, A., Dineen, D., Gibson, T. J., Karplus, K., Li, W., Lopez, R., Mcwilliam, H., Remmert, M., Soding, J., Thompson, J. D. & Higgins, D. G. 2011. Fast, scalable generation of high-quality protein multiple sequence alignments using Clustal Omega. Mol Syst Biol, 7, 539.

Simón-Soro, A., Tomás, I., Cabrera-Rubio, R., Catalan, M. D., Nyvad, B. & Mira, A. 2013. Microbial geography of the oral cavity. J Dent Res, 92, 616–21.

Spooner, R., Deguzman, J., Lee, K. L. & Yilmaz, O. 2014. Danger signal adenosine via adenosine 2a receptor stimulates growth of Porphyromonas gingivalis in primary gingival epithelial cells. Mol Oral Microbiol, 29, 67–78.

Spooner, R., Weigel, K. M., Harrison, P. L., Lee, K., Cangelosi, G. A. & Yilmaz, O. 2016. In Situ Anabolic Activity of Periodontal Pathogens Porphyromonas gingivalis and Filifactor alocis in Chronic Periodontitis. Sci Rep, 6, 33638.

Stark, H., Rodnina, M. V., Rinke-Appel, J., Brimacombe, R., Wintermeyer, W. & Van Heel, M. 1997. Visualization of elongation factor Tu on the Escherichia coli ribosome. Nature, 389, 403–6.

Su, Z., Kong, F., Wang, S., Chen, J., Yin, R., Zhou, C., Zhang, Y., He, Z., Shi, Y., Xue, Y., Shi, X., Lu, L., Shao, Q. & Xu, H. 2010. The rag locus of Porphyromonas gingivalis might arise from Bacteroides via horizontal gene transfer. Eur J Clin Microbiol Infect Dis, 29, 429–37.

Sun, Y., Wang, X., Li, J., Xue, F., Tang, F. & Dai, J. 2022. Extraintestinal pathogenic Escherichia coli utilizes the surface-expressed elongation factor Tu to bind and acquire iron from holo-transferrin. Virulence, 13, 698–713.

Szklarczyk, D., Gable, A. L., Lyon, D., Junge, A., Wyder, S., Huerta-Cepas, J., Simonovic, M., Doncheva, N. T., Morris, J. H., Bork, P., Jensen, L. J. & Mering, C. V. 2019. STRING v11: protein-protein association networks with increased coverage, supporting functional discovery in genome-wide experimental datasets. Nucleic Acids Res, 47, D607–D613.

Tamura, K., Stecher, G. & Kumar, S. 2021. MEGA11: Molecular Evolutionary Genetics Analysis Version 11. Mol Biol Evol, 38, 3022–3027.

Tomoyasu, T., Tabata, A., Imaki, H., Tsuruno, K., Miyazaki, A., Sonomoto, K., Whiley, R. A. & Nagamune, H. 2012. Role of Streptococcus intermedius DnaK chaperone system in stress tolerance and pathogenicity. Cell Stress Chaperones, 17, 41–55.

Torres, A. N., Chamorro-Veloso, N., Costa, P., Cadiz, L., Del Canto, F., Venegas, S. A., Lopez Nitsche, M., Coloma-Rivero, R. F., Montero, D. A. & Vidal, R. M. 2020. Deciphering Additional Roles for the EF-Tu, l-Asparaginase II and OmpT Proteins of Shiga Toxin-Producing Escherichia coli. Microorganisms, 8.

Tribble, G. D., Mao, S., James, C. E. & Lamont, R. J. 2006. A Porphyromonas gingivalis haloacid dehalogenase family phosphatase interacts with human phosphoproteins and is important for invasion. Proc Natl Acad Sci U S A, 103, 11027–32.

Tyanova, S., Temu, T. & Cox, J. 2016a. The MaxQuant computational platform for mass spectrometry-based shotgun proteomics. Nat Protoc, 11, 2301–2319.

Tyanova, S., Temu, T., Sinitcyn, P., Carlson, A., Hein, M. Y., Geiger, T., Mann, M. & Cox, J. 2016b. The Perseus computational platform for comprehensive analysis of (prote)omics data. Nat Methods, 13, 731–40.

Wellslager, B., Roberts, J., Chowdhury, N., Madan, L., Orellana, E. & Yilmaz, Ö. 2024. Porphyromonas gingivalis activates Heat-Shock-Protein 27 to drive a LC3C-specific probacterial form of select autophagy that is redox sensitive for intracellular bacterial survival in human gingival mucosa. bioRxiv, 2024.07.01.601539.

Whelan, S. & Goldman, N. 2001. A general empirical model of protein evolution derived from multiple protein families using a maximum-likelihood approach. Mol Biol Evol, 18, 691–9.

Widjaja, M., Harvey, K. L., Hagemann, L., Berry, I. J., Jarocki, V. M., Raymond, B. B. A., Tacchi, J. L., Gründel, A., Steele, J. R., Padula, M. P., Charles, I. G., Dumke, R. & Djordjevic, S. P. 2017. Elongation factor Tu is a multifunctional and processed moonlighting protein. Scientific Reports, 7, 11227.

Willis, J. R. & Gabaldón, T. 2020. The Human Oral Microbiome in Health and Disease: From Sequences to Ecosystems. Microorganisms, 8, 308.

Wilson, M. 2004. Microbial Inhabitants of Humans: Their Ecology and Role in Health and Disease, Cambridge, Cambridge University Press.

Wylensek, D., Hitch, T. C. A., Riedel, T., Afrizal, A., Kumar, N., Wortmann, E., Liu, T., Devendran, S., Lesker, T. R., Hernandez, S. B., Heine, V., Buhl, E. M., P, M. D. A., Cumbo, F., Fischoder, T., Wyschkon, M., Looft, T., Parreira, V. R., Abt, B., Doden, H. L., Ly, L., Alves, J. M. P., Reichlin, M., Flisikowski, K., Suarez, L. N., Neumann, A. P., Suen, G., De Wouters, T., Rohn, S., Lagkouvardos, I., Allen-Vercoe, E., Sproer, C., Bunk, B., Taverne-Thiele, A. J., Giesbers, M., Wells, J. M., Neuhaus, K., Schnieke, A., Cava, F., Segata, N., Elling, L., Strowig, T., Ridlon, J. M., Gulder, T. A. M., Overmann, J. & Clavel, T. 2020. A collection of bacterial isolates from the pig intestine reveals functional and taxonomic diversity. Nat Commun, 11, 6389.

Yang, J., Anishchenko, I., Park, H., Peng, Z., Ovchinnikov, S. & Baker, D. 2020. Improved protein structure prediction using predicted interresidue orientations. Proc Natl Acad Sci U S A, 117, 1496–1503.

Yao, L., Jermanus, C., Barbetta, B., Choi, C., Verbeke, P., Ojcius, D. M. & Yilmaz, O. 2010. Porphyromonas gingivalis infection sequesters pro-apoptotic Bad through Akt in primary gingival epithelial cells. Mol Oral Microbiol, 25, 89–101.

Yikilmaz, E., Chapman, S. J., Schrader, J. M. & Uhlenbeck, O. C. 2014. The interface between Escherichia coli elongation factor Tu and aminoacyl-tRNA. Biochemistry, 53, 5710–20.

Yilmaz, Ö. 2008. The chronicles of Porphyromonas gingivalis: the microbium, the human oral epithelium and their interplay. Microbiology (Reading), 154, 2897–2903.

Yilmaz, O., Verbeke, P., Lamont, R. J. & Ojcius, D. M. 2006. Intercellular spreading of Porphyromonas gingivalis infection in primary gingival epithelial cells. Infect Immun, 74, 703–10.

Yilmaz, O., Yao, L., Maeda, K., Rose, T. M., Lewis, E. L., Duman, M., Lamont, R. J. & Ojcius, D. M. 2008. ATP scavenging by the intracellular pathogen Porphyromonas gingivalis inhibits P2X7-mediated host-cell apoptosis. Cellular microbiology, 10, 863–875.

Yoshida, A., Nakano, Y., Yamashita, Y., Oho, T., Shibata, Y., Ohishi, M. & Koga, T. 1999. A novel dnaK operon from Porphyromonas gingivalis. FEBS Lett, 446, 287–91.

Zhang, Z., Liu, D., Liu, S., Zhang, S. & Pan, Y. 2021. The Role of Porphyromonas gingivalis Outer Membrane Vesicles in Periodontal Disease and Related Systemic Diseases. Frontiers in Cellular and Infection Microbiology, 10.

